# A Peroxiredoxin-P38 MAPK scaffold increases MAPK activity by MAP3K-independent mechanisms

**DOI:** 10.1101/2022.11.15.513554

**Authors:** Min Cao, Alison M Day, Martin Galler, Heather Latimer, Dominic P Byrne, Emilia Dwyer, Elise Bennett, Patrick A Eyers, Elizabeth A Veal

## Abstract

Peroxiredoxins (Prdx) utilize reversibly oxidized cysteine residues to reduce peroxides but also to promote H_2_O_2_ signal transduction, including H_2_O_2_-induced activation of P38 MAPK. Prdx form H_2_O_2_-induced disulfide complexes with many proteins, including multiple kinases involved in P38 MAPK signaling. Here we show that a genetically-encoded fusion between Prdx and the P38 MAPK is sufficient to hyperactivate the kinase in yeast and human cells by a mechanism that does not require the H_2_O_2_-sensing cysteine of the Prdx. In yeast, we demonstrate that a P38-Prdx fusion protein compensates for the loss of a scaffold protein and upstream MAP3K kinase activity, driving entry into mitosis. Based on our findings, we propose that the H_2_O_2_-induced formation of Prdx-MAPK disulfide complexes provides a scaffold and signaling platform for MAPKK-MAPK signaling. The demonstration that formation of a complex with a Prdx can be sufficient to modify the activity of a kinase has broad implications for peroxide-based signal transduction in eukaryotes.

**Highlights:** P38-Prdx complexes increase P38 (Sty1/MAPK14) phosphorylation in yeast and human cells

The *S. pombe* Prdx promotes transient thioredoxin-mediated oxidation of a MAPK tyrosine phosphatase

P38-Prdx complexes increase P38(Sty1) activity by phosphatase and MAP3K-independent mechanisms

P38-Prdx complexes increase the stability and phosphorylation of the *S. pombe* P38 MAPKK (Wis1)

Non-canonical, H_2_O_2_-induced autophosphorylation contributes to activation of the Wis1 MAPKK

## Introduction

Cells have evolved an array of sensing mechanisms to protect against damaging reactive oxygen species. These include redox-active peroxiredoxins, which form a key part of the cellular defence against peroxides and play important roles in aging and cancer (For a review see Nyström et al., 2012). The thioredoxin peroxidase activity of 2-Cys peroxiredoxins (Prdx) involves oxidation of the ‘peroxidatic’ cysteine-thiolate to sulfenate (SO^-^), followed by formation of a disulfide with a second ‘resolving’ cysteine in a neighboring peroxiredoxin. This Prdx-Prdx disulfide bond is subsequently reduced by thioredoxin (Trx)/thioredoxin reductase (TR) to complete the catalytic cycle (**Fig. 1A**) (For a review see (Bolduc et al., 2021). Thus, peroxiredoxins reduce peroxides. However, peroxiredoxins also have other roles in protecting cells against oxidative damage. For example, oligomeric forms of Prdx can act as chaperones, inhibiting the aggregation of proteins under stress conditions (Jang et al., 2004). As described below, Prdx also have important roles in promoting cell signaling in response to hydrogen peroxide (H_2_O_2_).

**Figure 1:**
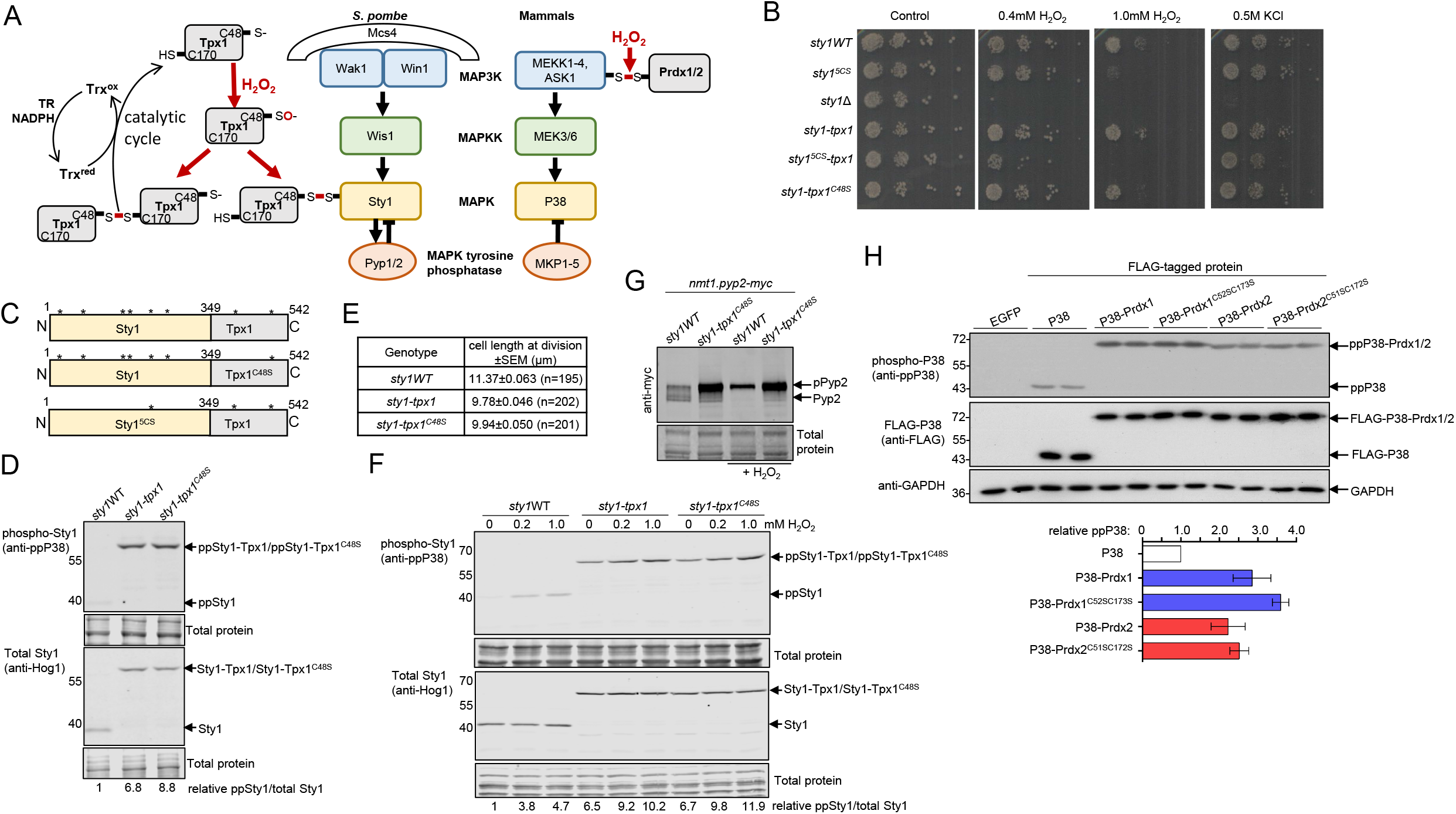
Constitutive formation of complex with a Prdx increases phosphorylation and activity of P38 MAPK in yeast and human cells. **[A]** Diagram illustrating (**left hand side**) the catalytic cycle of Prdx, exemplified by *S. pombe* Tpx1, which involves the sulfenylated (SO-) peroxidatic cys48 forming a disulfide bond with a resolving cys170 in another Tpx1 that is reduced by thioredoxin. Thioredoxin is then reduced by thioredoxin reductase(TR) using electrons from NADPH and (**right hand side**) the H_2_O_2_-induced formation of disulfide complexes between Prdx and components of P38 MAPK signaling pathways e.g. Sty1(P38). As illustrated, key components regulating activity of the *S. pombe* Sty1 and mammalian P38 MAPK by phosphorylation (MAPK kinase kinases (MAP3K), MAPK kinases (MAPKK) and MAPK tyrosine phosphatases (MKP) are conserved. Mcs4 is required for MAP3K activity in *S. pombe*. **[B]** Growth of equal numbers of exponentially growing *sty1WT(AD38), sty1^5CS^* (AD84), Δ*sty1* (JM1160), *sty1-tpx1* (MG17), *sty1^5CS^-tpx1* (MG23), and *sty1-tpx1^C48S^* (MG18) cells serially diluted and spotted on to YE5S plates containing H_2_O_2_, or 0.5M KCl, and incubated at 30°C for 3 days. (N=3) **[C]** Diagram of Sty1-Tpx1 fusion protein constructs. * indicates cysteine residues present at 13, 153, 158, 202, 242 in wild type Sty1 and residue 48 and 170 in wild-type Tpx1. In Sty1^5CS^ only C202 remains with the other 5 cysteines replaced by serine **[D]** The levels of Sty1 phosphorylation were determined by immunoblot analysis of cell lysates prepared from *sty1*WT(AD38), *sty1-tpx1*(MG17) and *sty1-tpx1^C48S^*(MG18) with antibodies against the dual TGY motif phosphorylated P38 (anti-ppP38); Total Sty1 levels were determined using anti-Hog1 antibodies. Total protein stain indicates protein loading. The positions of MW markers are shown (kD). The relative levels of phosphorylated Sty1/total Sty1 in each lane are shown **[E]** Length of exponentially growing cells at cell division. Cells expressing *sty1-tpx1* (MG17) or *sty1-tpx1^C48S^* (MG18) were significantly shorter than *sty1* WT (AD38) cells with P value<0.0001 (*t*-test) n=number of cells in each group N=2 **[F]** Sty1-phosphorylation was analysed by immunoblotting of lysate from exponentially growing WT (AD38) and *sty1-tpx1* (MG17) expressing cells before and following 10min exposure to the indicated concentrations of H_2_O_2_. Levels of Sty1 phosphorylation were normalised to total sty1 and relative levels are shown below lanes. **[G]** The mobility and levels of the Sty1-stabilized substrate, Pyp2, were examined in cells expressing Pyp2-Myc from the Sty1-independent *nmt1* promoter in *sty1WT* (EB15) or *sty1-tpx1^C48S^* (EB16) before and after 30 min exposure to 1.0 mM H_2_O_2_. N=3 **[H]** Phosphorylation of P38 in HEK-293T cells ectopically expressing EGFP (control), Flag epitope-tagged P38, P38-Prdx1, P38-Prdx2 or P38-Prdx fusion proteins in which the indicated cysteines in Prdx1 or 2 are serine substituted. The graph shows the mean relative phosphorylation of each protein compared with total protein (anti-Flag) determined from the 2 biological repeats shown. N=3. See also Figure S1 and Figure S2.

It is now well-established that hydrogen peroxide (H_2_O_2_) acts as a signaling molecule, initiating responses that protect against oxidative damage. H_2_O_2_ signals also regulate fundamental processes such as cell growth, differentiation or migration (for reviews see Holmstrom and Finkel, 2014; Hurd et al., 2012; Veal et al., 2007). Critical to the specificity of H_2_O_2_’s signaling function is its capacity to selectively promote the oxidation of specific protein-cysteine thiols and thus regulate the activity of targeted proteins, which include transcription factors (For example, Delaunay et al., 2002; Dansen et al., 2009), protein tyrosine phosphatases (for a review see Tonks, 2005) and kinases (For example, Byrne et al., 2020; Hourihan et al., 2016; Wani et al., 2011). However, Prdx are so abundant that H_2_O_2_ is unlikely to react with cysteines in these target proteins, before encountering the more highly reactive peroxidatic cysteine of a Prdx (Winterbourn, 2008). One solution to this problem, is that the sulfenylated Prdx, Prdx-SO^-^, formed following reaction of Prdx’s peroxidatic cysteine with H_2_O_2_, can act as the oxidant, forming a disulfide with a cysteine of a peroxide-targeted protein. This is exemplified by the H_2_O_2_-regulated transcription factors, STAT3, in mammalian cells, and AP-1-like transcription factors in yeast, where the formation of a transient disulfide complex with the peroxidatic cysteine of a Prdx initiates further oxidation events that regulate their activity (Delaunay et al., 2002; Okazaki et al., 2005; Sobotta et al., 2015; Bersweiler et al., 2017). This signaling function does not require the resolving cysteine and thioredoxin peroxidase activity of the Prdx. However, the thioredoxin peroxidase activity of Tpx1, the single 2-Cys Prdx and homolog of mammalian Prdx1 and Prdx2 in the fission yeast, *Schizosaccharomyces pombe*, is also essential for H_2_O_2_–induced activation of the AP-1-like transcription factor Pap1 (Bozonet et al., 2005; Brown et al., 2013; Vivancos et al., 2005). This is because thioredoxin reductase activity is limiting. Hence, the catalytic cycling of Prdx/Tpx1 as it reduces H_2_O_2_, also promotes the oxidation of thioredoxin family proteins, which reduce protein disulfides in many other proteins, including the intramolecular disulfides in active Pap1 (Day et al., 2012; Brown *et al*., 2013). Consequently, as abundant thioredoxin substrates, Prdx also act as H_2_O_2_-dependent inhibitors of thioredoxin’s oxidoreductase activity, thus regulating other enzymes that require reduced thioredoxin for their activity (Bodvard et al., 2017; Dangoor et al., 2012; Day et al., 2012; Ojeda et al., 2018).

P38 MAPKs are activated in response to a variety of different stimuli, including H_2_O_2_ (For reviews see Canovas and Nebreda, 2021; Cuenda and Rousseau, 2007). Canonical mechanisms of P38 activation involve regulation of both kinases and phosphatases, which together serve to modulate phosphorylation within the TGY motif in the kinase activation segment (**Fig. 1A**). The direct regulation of ‘upstream’ MAPKKs, alongside autophosphorylation and Arg-methylation based mechanisms of P38 itself, have been proposed to control the signaling outputs of P38 (Johnson and Lapadat, 2002; Sanz-Ezquerro and Cuenda, 2021; Canovas and Nebreda, 2021). The *S. pombe* P38-related Sty1 MAPK (also known as Spc1) is activated by a wide range of environmental stimuli (Millar et al., 1995; Shiozaki and Russell, 1995; Degols et al., 1996). Interestingly, the *S. pombe* Prdx, Tpx1, is specifically required for the activation of Sty1/P38 in response to H_2_O_2_ (Veal et al., 2004). Prdx have also been shown to be important for activating P38 MAPK in metazoa (Barata and Dick, 2020; Conway and Kinter, 2006; De Haes et al., 2014; Jarvis et al., 2012; Olahova et al., 2008). In mammals, the Prdx, Prdx1, has been proposed to instigate P38 activation by promoting the disulfide bond-mediated oligomerisation and activation of the ‘upstream’ Ser/Thr MAP kinase kinase kinase (MAP3K), Ask1 (Jarvis *et al*., 2012). Intermolecular disulfides formed between Prdx2 and MAP3Ks have also been implicated in the activation of P38 in human and Drosophila cells (Barata and Dick, 2020). Intriguingly, H_2_O_2_-induced disulfides also form between the peroxidatic cysteine in Tpx1 and cysteines in Sty1 (Veal *et al*., 2004). However, the role that Tpx1-Sty1 disulfide complexes play in promoting Sty1 (p38) phosphorylation has remained unclear: Indeed, despite intense scrutiny, we have been unable to find any evidence that Sty1-Tpx1 disulfides are an intermediate in the generation of intramolecular or intermolecular disulfides between cysteines in Sty1, as observed for Yap1, STAT3 and ASK1 (Delaunay *et al*., 2002; Jarvis *et al*., 2012; Sobotta *et al*., 2015).

In this paper, we have tested an alternative hypothesis; that formation of a complex with Tpx1/Prdx might be sufficient to increase Sty1/P38 phosphorylation. To test this we have undertaken a genetically-encoded proximity fusion approach, similar to those used to evaluate the function of covalently linked interactions with ubiquitin-like proteins, such as SUMO (Ross et al., 2002), and the effect of physical constraints on signaling outputs from the PKA holoenzyme complex (Smith et al., 2017).

Using this approach, we reveal that formation of a constitutive complex with a Prdx is sufficient to activate P38 in both yeast and human cells, even in the absence of any activating stimuli. We demonstrate that Prdx-P38 complexes activate the *S. pombe* P38, Sty1, by promoting both the canonical and non-canonical phosphorylation of the MAPKK. Notably, we find that the peroxide-reacting cysteine of the Prdx is dispensable for the increased activity of P38-Prdx complexes in driving entry into mitosis. Instead, our data support a model in which the Prdx-MAPK fusion protein provides a scaffold for MAPKK-MAPK signaling. In addition, we show that increased levels of Prdx (Tpx1) stimulate the H_2_O_2_-induced formation of disulfide complexes involving thioredoxin (Trx1) and the MAPK phosphatase, Pyp1. This suggests that the thioredoxin peroxidase activity of Tpx1/Prdx may also increase H_2_O_2_-induced Sty1/P38 phosphorylation by promoting the oxidation of thioredoxin and the MAPK phosphatase Pyp1. We propose that together these complementary mechanisms allow fission yeast to ‘stock-take’ their reductive capacity to tailor MAPK-dependent responses, and the timing of entry into mitosis, according to the level of oxidative stress.

## Results

### Multiple cysteines in Sty1(P38) are involved in Tpx1-Sty1 disulfides and important for Sty1 activity

Following exposure of cells to H_2_O_2_, disulfide complexes form between the invariant peroxidatic residue(C48) of the Prdx, Tpx1 and the P38 MAPK, Sty1 (**Fig. 1A**) (Veal *et al*., 2004). Our previous work identified that cysteine 35 in Sty1 was important for formation of one of the disulfide complexes with Tpx1 (Veal et al 2004). However, Tpx1-Sty1 disulfide complexes still form in cells expressing Sty1^C35S^ ectopically (Veal et al 2004) or from the *sty1* chromosomal locus (**Fig. S1A**). Moreover, overexpression of Tpx1 still stimulates increased H_2_O_2_-induced phosphorylation of Sty1^C35S^ on the key regulatory Thr/Tyr residues in the activation segment, indicating that the effect of Tpx1 is not solely dependent on C35 (**Fig. S1B**). Intriguingly, five of Sty1’s six cysteines are predicted to be on the surface of Sty1 where they would be available to react with H_2_O_2_/Prdx (Tpx1), with only C202 internal and required for Sty1 stability (Day and Veal, 2010). The conservation of several of these surface cysteines in other MAPK is consistent with data suggesting they are redox-sensitive and important for function (Day and Veal, 2010; Marino and Gladyshev, 2010; Veal *et al*., 2004). Indeed, our further analysis has revealed that only in cells expressing a Sty1 mutant in which these five cysteines were all substituted with serine (Sty1^5CS^), was Sty1-Tpx1 disulfide formation abrogated **(Fig. S1C)**. Thus, consistent with our earlier work, multiple cysteines in Sty1 are able to form disulfides with Tpx1 (Veal et al 2004).

This oxidation-resistant Sty1^5CS^ mutant provided an opportunity to examine the role of Tpx1-Sty1 disulfides in regulating Sty1 phosphorylation. However, our analysis of Sty1^5CS^-mutant expressing cells suggested that this Sty1 mutant protein was only partially functional: Sty1 activity is important for determining the timing of cell entry into mitosis with Δ*sty1* cells significantly longer that wild-type cells, as a result of a longer G2 phase. Conversely, increased Sty1 activity shortens G2 phase such that cells divide at a smaller size (Millar et al., 1995; Shiozaki and Russell, 1995). Our analysis revealed that Sty1^5CS^-mutant expressing cells were larger and divided at an increased length compared with wild-type cells (**Fig. S1D-E**). This was despite slightly elevated levels of Sty1^5CS^ phosphorylation compared with wildtype Sty1 (**Fig. S1F**). Sty1 is also important for adaptation/survival under a number of different stress conditions, including osmotic and oxidative stress (Millar et al., 1995; Shiozaki and Russell, 1995)(**Fig. 1B**). Sty1^5CS^-expressing cells were able to adapt to osmotic stress, but, as expected from previous work (Day and Veal, 2010), had a reduced oxidative stress tolerance, particularly at higher levels of H_2_O_2_ (**Fig. 1B**). Intriguingly, we observed that Sty1^5CS^-expressing cells contained lower levels of the MAPK phosphatase, Pyp1, than wildtype cells (**Fig. S1G**) providing a possible explanation for the slightly increased phosphorylation of Sty1^5CS^ (**Fig. S1F**). Together these data suggest that cysteines involved in Sty1-Tpx1 disulfides are important for Sty1 activity under physiological and oxidative stress conditions and may contribute to a negative feedback mechanism regulating Pyp1.

### Constitutive formation of complex with a Prdx increases phosphorylation of *S. pombe* Sty1 and human P38 MAPK

Next we evaluated whether constitutive formation of a complex between the Prdx, Tpx1, and Sty1 (P38) was sufficient to activate Sty1. To do this we engineered cells to express Sty1-Tpx1 fusion proteins from the Sty1 chromosomal locus **(Fig. 1C)**. Overexpression of Tpx1, increases Sty1 activation in response to H_2_O_2_ (**Fig. S1B**) by a mechanism that requires the peroxide-reacting cysteine 48 of Tpx1 (Veal *et al*., 2004). Therefore, it was possible that any effect of a Tpx1 fusion could be an indirect consequence of increasing levels of active Prdx (Tpx1). Hence, to eliminate this possibility, we also generated cells expressing a Sty1-Tpx1^C48S^ fusion protein, in which the peroxide-reacting cysteine, that is also engaged in Tpx1-Sty1 disulfides (Veal *et al*., 2004) (**Fig. 1A**), was substituted with serine. It was possible that the juxtaposition of the two proteins might cause issues with their folding. However, both Sty1-Tpx1 fusions were expressed at similar levels to wild-type Sty1 **(Fig. 1D)** and, importantly, both supported growth under control and stress conditions **(Fig. 1B)**. This suggested that Sty1 function was retained in both Sty1-Tpx1 fusion proteins. In contrast, fusion of Tpx1 to the Sty1^5CS^ mutant did not rescue the oxidative stress sensitivity associated with expression of Sty1^5CS^ (**Fig. 1B**). This is consistent with previous work indicating that cysteines in Sty1 contribute to Sty1 function independently from their role in Tpx1-Sty1 disulfide complex formation (**Fig. 1B, S1D-E**) (Day and Veal, 2010).

Notably, we observed increased Sty1 phosphorylation in cells expressing Sty1-Tpx1 or Sty1-Tpx1^C48S^, even in the absence of stress **(Fig. 1D)**. This was consistent with our hypothesis that the interaction with Tpx1 was sufficient to increase Sty1 activity. However, increased Sty1 phosphorylation was also observed in Sty1^5CS^-expressing cells in which Sty1 function is impaired (**Fig. 1B, S1D-F**). Hence, it was also possible that increased phosphorylation of the Sty1-Tpx1 fusion proteins reflected impaired Sty1 activity and, consequently, reduced feedback inhibition by Sty1-regulated MAPK phosphatases, Pyp1 and Pyp2 (Kowalczyk *et al*., 2013) (**Fig. 1A, S1G**). However, in contrast to Sty1^5CS^-expressing cells, there was no reduction in the levels of Pyp1 in cells expressing Sty1-Tpx1 fusion proteins (**Fig. S1G**). Nevertheless, it was important to determine whether the increased Sty1 phosphorylation correlated with increased Sty1 activity. Increased Sty1 catalytic activity is associated with reduced cell size at division, resulting from an accelerated entry into mitosis (Millar *et al*., 1995; Shiozaki and Russell, 1995). Therefore, to investigate whether the increased phosphorylation of Sty1-Tpx1 fusion proteins **(Fig. 1D)** was associated with increased Sty1 activity, we examined the length at which cells expressing wild-type or Sty1-Tpx1 fusion proteins divided. This revealed that Sty1-Tpx1 and Sty1-Tpx1^C48S^ expressing cells divided at a significantly smaller size than wild type cells. Importantly, this indicated that constitutive formation of a complex with Tpx1 increased both the phosphorylation and activity of Sty1 **(Fig. 1E)**. Notably, the level of phosphorylation of the Sty1-Tpx1 fusion proteins under normal growth conditions was similar to the level of Sty1 phosphorylation induced in wild-type cells following exposure to levels of H_2_O_2_ that stimulate Sty1-Tpx1 disulfide formation **(Fig. 1F, S1A, C)**(Veal et al., 2004).

We also examined how expression of a Sty1-Tpx1^C48S^ fusion affected the phosphorylation of a Sty1 substrate, the MAPK-phosphatase Pyp2, that is stabilized by Sty1-dependent phosphorylation (Kowalczyk *et al*., 2013). As expected, and consistent with Sty1-dependent phosphorylation, H_2_O_2_-treatment was required to shift the mobility of Pyp2 (expressed from the Sty1-independent *nmt1* promoter) in wild-type cells (**Fig. 1G**). In contrast, cells expressing Sty1-Tpx1^C48S^ contained substantially increased levels of lower mobility phosphorylated Pyp2 (p-Pyp2), even prior the addition of H_2_O_2_. These data strongly support the conclusion that the Sty1-Tpx1^C48S^ fusion protein is constitutively hyperactive compared with wild-type Sty1 **(Fig. 1E and 1G)**.

Prdx have also been shown to stimulate activation of P38 MAPK in animals (Conway and Kinter, 2006; Jarvis *et al*., 2012; Olahova *et al*., 2008). Moreover, although P38-Prdx disulfides have not been reported in mammalian cells, Prdx have been shown to form disulfide complexes with kinases upstream of P38 (Barata and Dick, 2020; Jarvis *et al*., 2012)(**Fig. 1A**). Thus, we tested whether constitutive fusion to the cytosolic Prdx1 or Prdx2 was able to activate P38 (MAPK14) in human cells. Importantly, P38-Prdx1 and P38-Prdx2 fusions were both expressed at similar levels to P38 and demonstrated catalytic activity towards a model P38 peptide after immunoprecipitation (**Fig. 1H and S2**). Although, as expected, phosphorylation of P38 was very low under normal culture conditions, the phosphorylation of both P38-Prdx1 and P38-Prdx2 fusions was significantly increased (**Fig. 1H**). This indicated that formation of a complex with a Prdx is also sufficient to increase the phosphorylation of human P38. Moreover, similar to our findings with Sty1-Tpx1^C48S^, neither the peroxide-reacting or resolving cysteine of Prdx1 or Prdx2 were required for enhanced P38 phosphorylation (**Fig. 1H**). Thus, we have established that the function of P38-Prdx complexes in stimulating increased P38 phosphorylation at the key regulatory Thr/Tyr residues in the activation segment is conserved.

### Tpx1 (Prdx) promotes Sty1 (P38) activation downstream/independently of MAP3K and a redox-sensitive cysteine in the MAPKK, Wis1

Next, we investigated the mechanism(s) by which P38-Prdx complexes increase P38 activity. Taking advantage of the wealth of genetic tools available in *S. pombe*, we focused on identifying how Sty1-Tpx1 complexes activated Sty1. In contrast with the redundancy of mammalian MKK3 and MKK6, a single MAPKK, Wis1, is responsible for phosphorylating Sty1 (Millar *et al*., 1995; Shiozaki and Russell, 1995). Interestingly, a cysteine in Wis1, C458, was recently proposed to be sensitive to oxidation and involved in regulating Wis1 activity (Sjolander et al., 2020). Thus, it was possible that C458 was important for the effect of Tpx1 on Sty1 activation. However, ectopic overexpression of Tpx1 (**Fig. S3A**) and expression of Sty1-Tpx1 fusion protein (**Fig. S3B**) both caused similar increases in Sty1 phosphorylation in cells expressing Wis1^C458S^ or wild-type Wis1. Thus, it was unlikely that altered oxidation of C458 mediated the effect of Tpx1 on Sty1 phosphorylation (**Fig. S3A-B**).

Our previous work demonstrated that overexpression of Tpx1 restores peroxide-inducible Sty1 phosphorylation to cells expressing a phospho-mimetic Wis1 mutant (Wis1^DD^), in which the MAP3K-phosphorylated sites (Ser469 and Thr473) are substituted with aspartate (Veal *et al*., 2004). This suggests that overexpression of Tpx1 does not stimulate H_2_O_2_-dependent Sty1 phosphorylation by increasing MAP3K-dependent activation of Wis1. Moreover, when we attempted to construct a strain co-expressing Sty1-Tpx1 and the phospho-mimetic Wis1^DD^ mutant, less than 10% of viable spores obtained from a cross between the 2 strains bore markers that indicated the presence of both alleles (rather than the expected 25% for unlinked genes). Sty1 activity is tightly regulated such that Wis1 overexpression is lethal (Shiozaki and Russell, 1995). Hence, the reduced viability of cells co-expressing *sty1-tpx1* and *wis1^DD^* suggested that there might be a synthetic negative interaction, with both alleles acting independently to increase Sty1 phosphorylation to lethal levels. Indeed, microscopic examination and immunoblot analysis indicated that *‘sty1-tpx1 wis1DD’* strains, which genotyping indicated bore both alleles, had adapted to the deleterious effect of hyperactivated Sty1, by eliminating or reducing Sty1 expression to such an extent that they exhibited the long cell phenotype associated with its loss (**Fig. S3C-D**). Taken together these data strongly suggest that Sty1-Tpx1 complexes and overexpression of Tpx1 both increase phosphorylation of Sty1 by a novel MAP3K-independent mechanism(s).

### High levels of the Prdx, Tpx1, promote H_2_O_2_-induced Sty1 (P38) phosphorylation by increasing the thioredoxin-dependent oxidation of the MAPK tyrosine phosphatase Pyp1

Next we evaluated whether Tpx1 might increase H_2_O_2_-inducible Sty1 phosphorylation by inhibiting the dephosphorylation of Sty1. The protein tyrosine phosphatase Pyp1 is a MAPK phosphatase that is important for maintaining low levels of Sty1 activity under non-stress conditions, with the stress-induced MAPK phosphatase, Pyp2, also responsible for inhibition of Sty1 through a negative feedback mechanism (Millar *et al*., 1995) (**Fig. 1A, 2A-B**). The utilization of a deprotonated cysteine in the catalytic site of protein tyrosine phosphatases renders them highly sensitive to inactivation by peroxide (Tonks, 2005). Hence, we considered the possibility that Tpx1 increased H_2_O_2_-induced Sty1 activation by promoting the oxidation of Pyp1 or Pyp2. First we tested whether either Pyp1 or Pyp2 was required for the elevated H_2_O_2_-induced Sty1 phosphorylation in cells ectopically overexpressing Tpx1 (**Fig. 2A, S1B**) (Veal et al, 2004). As expected, in Δ*pyp1* mutant cells, Sty1 phosphorylation was increased, confirming the importance of this phosphatase in maintaining low levels of Sty1 phosphorylation (Millar *et al*., 1995; Shiozaki and Russell, 1995)(**Fig. 2A**). Moreover, there was no significant increase in Sty1 phosphorylation in response to H_2_O_2_, even in Δ*pyp1* cells overexpressing Tpx1 **(Fig. 2A-B)**. In contrast, as in wild-type cells (**Fig. S1B**, Veal et al 2004), overexpression of Tpx1 increased H_2_O_2_-induced Sty1 phosphorylation in cells expressing pk-tagged Pyp1, and also restored some H_2_O_2_-inducibility to Sty1 phosphorylation to Δ*pyp2* cells (**Fig. 2B**). This suggested that increased Tpx1 levels might promote increased H_2_O_2_-induced Sty1 phosphorylation by promoting the oxidation and inactivation of Pyp1. To test this hypothesis, we examined the effect of Tpx1 on Pyp1 oxidation. This revealed that Pyp1 was susceptible to transient oxidation to slower-migrating forms (**Fig. 2C**). These slower-migrating forms were absent when samples were pre-treated with the reducing agent 2-mercaptoethanol prior to SDS-PAGE, identifying them as disulfide-bonded complexes containing Pyp1 (see ‘Total Pyp1’ in lower panels of **Fig. 2C-D, 2F** and not shown). Notably, these complexes were not detected in Δ*tpx1* mutant cells, suggesting that Tpx1 is required for their formation (**Fig. 2C, 2E**). Moreover, the formation of these oxidized forms of Pyp1 was dramatically increased in cells expressing high levels of Tpx1, such that reduced Pyp1 levels were depleted to less than 50% of total Pyp1 **(Fig. 2D)**. These data suggest that Tpx1 promotes reversible peroxide-induced oxidation of Pyp1 into slower-migrating complexes. The correlation between H_2_O_2_-induced formation of these complexes and increased Sty1 phosphorylation is consistent with the formation of these complexes lowering Pyp1 activity towards Sty1 (**Fig. 2A-D**).

**Figure 2:**
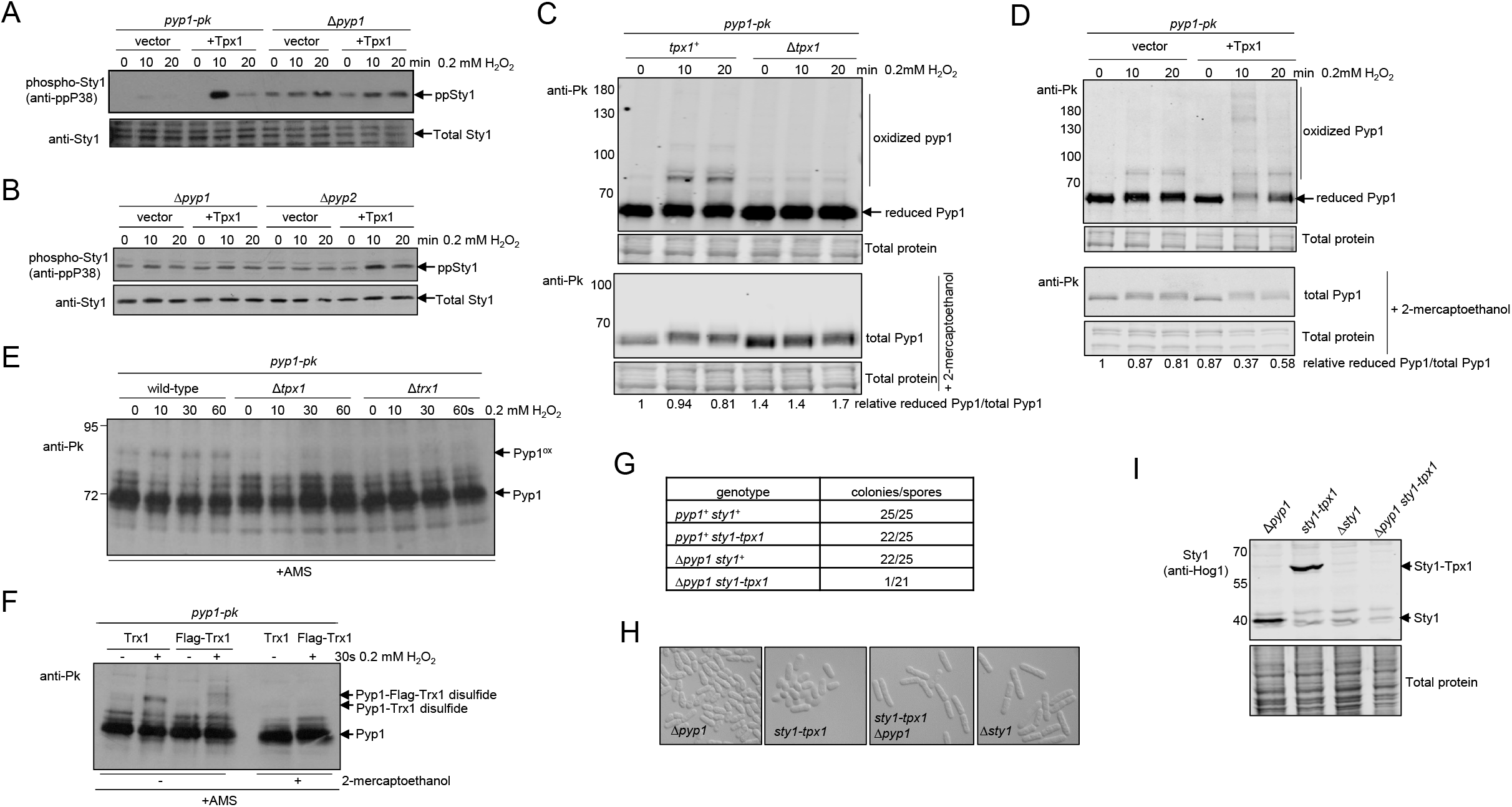
High levels of Prdx(Tpx1) promote H_2_O_2_-induced P38(Sty1) phosphorylation by promoting the thioredoxin-dependent oxidation of the MAPK tyrosine phosphatase Pyp1. Immunoblot analysis of **[A-B]** Sty1 phosphorylation in wild type cells expressing *pyp1-pk* (HL2), Δ*pyp1* cells (NJ102) and Δ*pyp2* (JP279) cells expressing vector (Rep1) or Rep1 *tpx1*^+^, before and following 10 or 20 min exposure to 0.2 mM H_2_O_2_. **[C-F]** Analysis of Pyp1 oxidation state in cells expressing Pk-tagged Pyp1 (Pyp1-3pk) that was detected with anti-Pk antibodies in **[C]** wild-type *tpx1*^+^(AD142) and Δ*tpx1* (AD143) cells. Samples reduced with 2-mercaptoethanol were used to determine ‘total Pyp1’. The relative levels of ‘reduced Pyp1’ normalised to ‘total Pyp1’ are shown. **[D]** wild-type *tpx1*^+^ cells (HL2) expressing vector (Rep1) or Rep1*tpx1*^+^ before and following treatment with 0.2 mM H_2_O_2_ for 0, 10 and 20 min. Samples treated with 2-mercaptoethanol were used to detect ‘total Pyp1’. The relative levels of ‘reduced Pyp1’ normalised to ‘total Pyp1’ are shown. **[E]** wild-type (AD142), Δ*tpx1* (AD143) and Δ*trx1* (AD130) cells before or after 10, 30 or 60s exposure to 0.2 mM H_2_O_2_. Proteins were reacted with AMS during solubilization step. The **[F]** wild-type Trx1 (AD142) or Flag-tagged Trx1 (AD144) before or after 30s exposure to 0.2 mM H_2_O_2_. Proteins were reacted with AMS during solubilization step. The sensitivity of the band with lower mobility in Flag-Trx1-expressing cells to 2-mercaptoethanol treatment confirming that it represents a Trx1-Pyp1 disulfide complex. **[G]** Analysis of dissected tetrads from a cross between Δ*pyp1 sty1^+^* (NJ102) and *pyp1^+^ sty1-tpx1* (MG17) cells showing the number of viable spores (colonies formed)/total number of spores of that genotype. **[H]** DIC light microscopy of exponentially growing Δ*pyp1* (NJ102), *sty1-tpx1* MG17, Δ*pyp1 sty1-tpx1* (MC150) cells in comparison with a Δ*sty1* (JM1160). **[I]** Immunoblotting analysis of Δ*pyp1 sty1*^+^ (NJ102) and *sty1-tpx1 pyp1*^+^ (MG17) Δ*sty1* (JM1160) and Δ*pyp1 sty1-tpx1* (MC150) with anti-Hog1 antibodies demonstrated that Sty1-Tpx1 fusion protein was undetectable in the single isolate of Δ*pyp1 sty1-tpx1* cells (MC150) obtained from crossing NJ102 and MG17. See also Figure S3.

Next, we explored the role Tpx1 plays in promoting the formation of these Pyp1-containing complexes. First, we determined that the most rapidly induced complex (~84kD) was detected within 10s following exposure of wild-type cells to 0.2 mM H_2_O_2_ **(Fig. 2E**). Thioredoxin family proteins form disulfide complexes with protein tyrosine phosphatases (PTPs) that have been proposed to be important for redox-regulation of PTP activity (Dagnell et al., 2013; Netto and Machado, 2022; Schwertassek et al., 2014). Indeed, we noted that the mobility (~84kD) of this H_2_O_2_-induced complex was consistent with that expected for a disulfide between Pyp1.3pk (72kD) and Trx1(11.3kD) (**Fig. 2E**) (Dagnell *et al*., 2013; Schwertassek *et al*., 2014). Thus, to investigate these complexes further, we examined them in a mutant lacking the thioredoxin, Trx1. This revealed that Trx1 was required for the formation of Pyp1 disulfide complexes (**Fig. 2E**). Moreover, in cells expressing FLAG epitope-tagged Trx1 instead of wild-type Trx1, the most prominent oxidized (2-mercaptoethanol-sensitive) Pyp1 species formed was reduced in mobility, consistent with it representing a disulfide complex between Pyp1 and Trx1 (**Fig. 2F**). Our data are consistent with the thioredoxin peroxidase activity of Tpx1 stimulating the formation of these complexes by promoting the oxidation of Trx1: In response to these low (0.2 mM) levels of H_2_O_2_ Tpx1-Tpx1 disulfides become the most abundant thioredoxin substrate and TR activity is limiting, such that the pool of Trx1 is transiently oxidized (Day *et al*., 2012). Therefore, increased Tpx1 activity would be expected to inhibit the reduction of disulfides in oxidized Pyp1 complexes, as we observe (**Fig. 2C-D**). Intriguingly, the very rapid formation of the Pyp1-Trx1 disulfide complex (**Fig. 2E-F**) also raises the possibility that, in these circumstances, where other reductants are limited, oxidized Trx1 can accept an electron from a Pyp1 cysteine/s to form a Pyp1-Trx1 disulfide. The nature of the other lower migrating Pyp1 forms remains to be established, and further work would be required to establish how these disulfide complexes regulate Pyp1 activity. However, Pyp1 is known to be susceptible to aggregation under stress conditions, with this aggregation resulting in increased Sty1 activation under these conditions (Boronat et al., 2020; Nguyen and Shiozaki, 1999). Hence, given their association with increased Sty1 phosphorylation (**Fig. 2**), it is possible that formation of the disulfides we detect may provide a reversible mechanism to transiently inhibit Pyp1.

### Sty1-Tpx1 (P38-Prdx) complexes increase Sty1 activity independently from H_2_O_2_/Tpx1/Trx1-dependent regulation of Pyp1

Loss of *pyp1* and expression of Sty1-Tpx1 or Sty1-Tpx1^C48S^ fusion proteins both increase Sty1 phosphorylation (**Figs. 1D, 1F, 2A, 2B**). Sty1-Tpx1 fusion proteins also increase the cellular Tpx1 (Prdx) content. Thus, it was possible that, similar to overexpressing Tpx1, Sty1-Tpx1 (P38-Prdx) complexes also activated Sty1(P38) by a *pyp1*-dependent mechanism. To test this hypothesis we crossed strains expressing Sty1-Tpx1 fusion and Δ*pyp1* mutant alleles. Our expectation was that, if Sty1-Tpx1 fusions act solely via inhibiting Pyp1, there should be no further increase in Sty1 phosphorylation in cells expressing both alleles. However, instead we observed a very strong (95%) synthetic lethal interaction, suggesting that when both alleles are expressed in the same cell they increase Sty1 phosphorylation to lethal levels **(Fig. 2G** and not shown). Moreover, analysis of surviving, slow-growing colonies bearing markers of both alleles, revealed that these cells had completely eliminated Sty1-Tpx1 expression and activity **(Fig. 2H-I** and not shown). These results indicate a very strong selection pressure to eliminate Sty1-Tpx1 expression in Δ*pyp1* mutant cells. This indicates that the physiological impacts of constitutive Sty1 activation in cells expressing Sty1-Tpx1/Sty1-Tpx1^C48S^ fusion proteins are partially mitigated by Pyp1. Importantly, these data strongly suggest that Sty1-Tpx1 complex formation can independently increase Sty1 activity aside from any effect on Pyp1 oxidation state.

### High levels of H_2_O_2_ are required for maximal, MAP3K-dependent phosphorylation of the Wis1 MAPKK

The synthetic lethality of Sty1-Tpx1-expressing alleles with either a Δ*pyp1* deletion mutant or a *wis1DD* mutant allele (in which canonical MAP3K sites in the MAPKK are substituted with phospho-mimetic residues) (**Figs. 2G-I, S3C-D**) strongly suggested that Sty1-Tpx1 fusions increase Sty1 phosphorylation by novel MAP3K and phosphatase-independent mechanism/s. Wis1 is the only ‘upstream’ MAPKK that phosphorylates Sty1 (Millar *et al*., 1995; Shiozaki and Russell, 1995). Therefore, we examined the effect of Sty1-Tpx1 on activation of Wis1. First, we established an assay to evaluate the activation of Wis1 by phosphorylation. Using Phos-tag^TM^ to enhance the separation of phosphorylated forms, we detected lower mobility Wis1 and Wis1^DD^ species in cells grown under non-stress conditions that were eliminated by phosphatase treatment (**Fig. S4A**). This suggests that Wis1 is phosphorylated under physiological conditions, on different sites from those regulated by the MAP3K. Wis1 is required for increased Sty1 phosphorylation in response to various stresses, including osmotic and oxidative stress (Millar *et al*., 1995; Shiozaki and Russell, 1995). Indeed, as predicted, we detected a phosphorylation-dependent decrease in Wis1 mobility under osmotic stress conditions (**Fig. S4A**). As expected, given the stress-induced activation of the MAP3Ks, Wak1 and Win1 (**Fig. 1A**), we observed higher levels of phosphorylation of wild-type Wis1, which contains MAP3K-phosphorylated residues, than for the Wis1^DD^ mutant (**Fig. S4A**). However, when we examined the effect of increasing concentrations of H_2_O_2_ we were surprised to find that exposure to a much higher dose (≥1 mM) was required to produce detectable increases in Wis1 phosphorylation than the dose of H_2_O_2_ required to activate Sty1 (**Fig. 3A, 1F, S4B**). This suggested that the Tpx1-dependent oxidation of Pyp1 and formation of Tpx1-Sty1 disulfide complexes detected in response to 0.2 mM H_2_O_2_ (**Fig. S1A**, **Fig. 2C-F**) might be more important for activation of Sty1 by these lower levels (≤ 1 mM).

**Figure 3.**
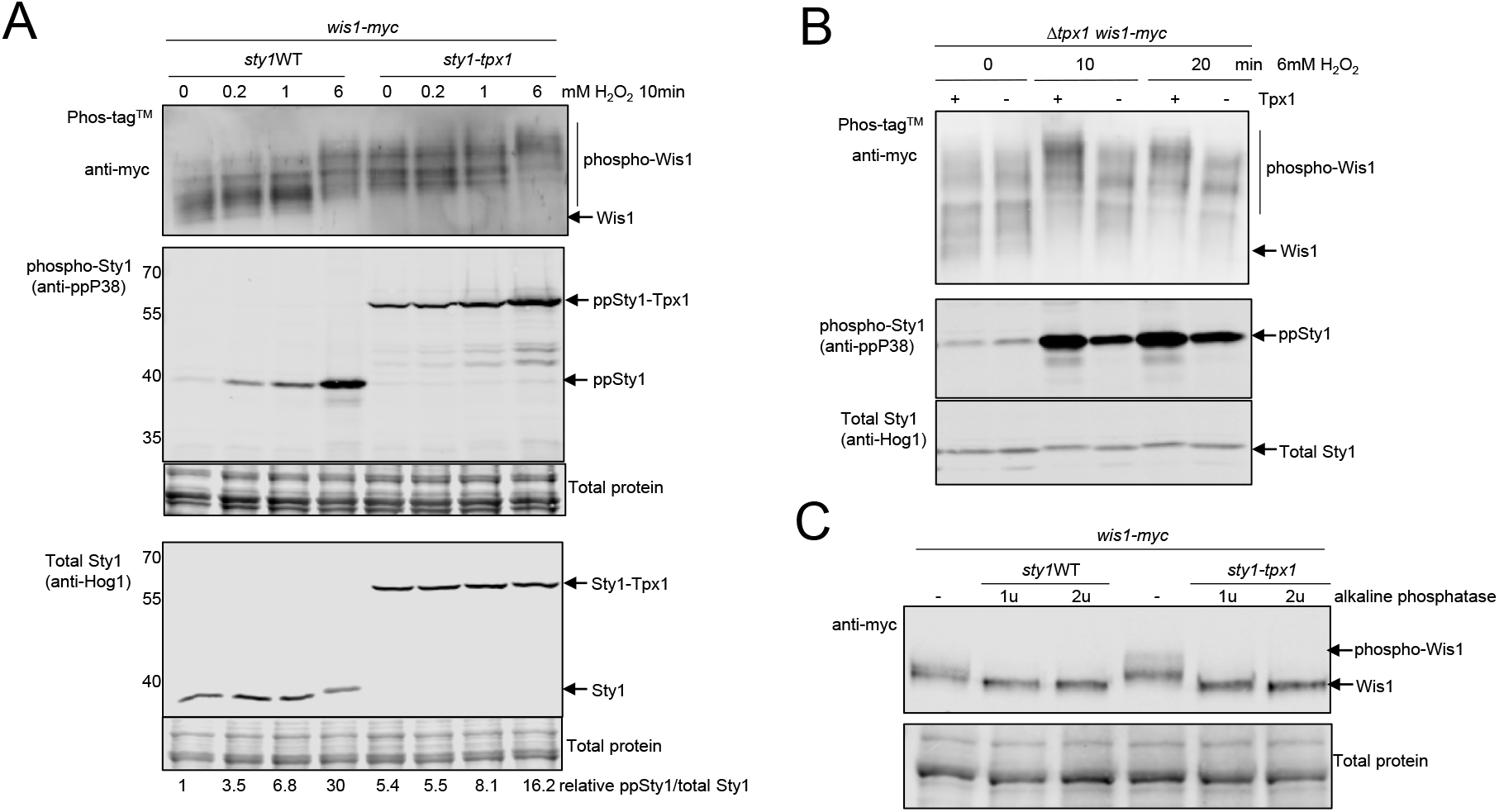
H_2_O_2_-induced phosphorylation of the MAPKK Wis1 requires Prdx(Tpx1) and P38-Prdx(Sty1-Tpx1) complexes promote constitutive Wis1 phosphorylation. The reduced mobility of phosphorylated Wis1 was detected by immunoblotting proteins separated using Phos-tag^TM^ SDS-PAGE. **[A]** Wis1 and Sty1 phosphorylation in wild type cells (MC2) and cells expressing Sty1-Tpx1 fusion protein (MC12) before and following exposure to increasing concentrations of H_2_O_2_. The relative levels of dual TGY motif phosphorylated Sty1 phosphorylation (anti-ppP38) normalised to total Sty1 (anti-Hog1) are shown below each lane. **[B]** The increased phosphorylation of Wis1 and Sty1 in response to 6 mM H_2_O_2_ [A] was impaired in Δ*tpx1* delete cells (MC88) containing vector (Rep1) but restored by expression of Tpx1 from *Rep1tpx1^+^*. The levels of dual TGY motif phosphorylated Sty1 phosphorylation (anti-ppP38) normalised to total Sty1 (anti-Hog1) are also shown. **[C]** Comparison of the mobility of Wis1 in untreated and alkaline phosphatase-treated extracts from cells expressing wild-type Sty1 (MC2) or *sty1-tpx1* (MC12). The phosphatase-sensitivity of the less mobile forms is consistent with the mobility shift in Wis1 reflecting increased phosphorylation. The positions of MW markers separated on conventional gels are indicated (kD). See also Figure S4.

Intriguingly, we also observed that Wis1^DD^ was phosphorylated following stress, despite the absence of the canonical phosphosites for the upstream MAP3K (**Fig. S4A, S4C**). However, importantly, Wis1^DD^ mobility was affected to a much lesser extent in response to either stress, indicating that the more pronounced shift in wild-type Wis1 was due to phosphorylation of canonical MAP3K-phosphorylated residues (**Fig. S4C**). As expected, H_2_O_2_-induced Wis1 phosphorylation was dependent on the presence of Mcs4, a fungal-specific scaffold protein required to support the activity of the MAP3Ks, Win1 and Wak1 (**Fig. S4D, Fig. 1A**) (Morigasaki and Shiozaki, 2013; Shiozaki et al., 1997). Having established that Wis1 undergoes multiple phosphorylation events, with phosphorylation on MAP3K sites only increasing significantly following exposure of cells to higher concentrations of H_2_O_2_ (6 mM), next we investigated whether Tpx1-Sty1 complexes affected Wis1 phosphorylation.

### Tpx1 is required for H_2_O_2_-induced Wis1 phosphorylation and Sty1-Tpx1 complexes increase Wis1 phosphorylation

Consistent with the specific requirement for Tpx1 in H_2_O_2_-induced activation of Sty1, over a range of concentration, up to 10 mM H_2_O_2_ (Veal *et al*., 2004), analysis of Δ*tpx1* mutant cells indicated that Tpx1 was important for Wis1 phosphorylation in response to H_2_O_2_ but not osmotic stress (**Fig. 3B, S4E-F**). Next, we examined whether Wis1 phosphorylation was affected in Sty1-Tpx1 fusion expressing cells. Strikingly, we observed that Wis1 underwent a shift consistent with phosphorylation in cells expressing Sty1-Tpx1 or Sty1-Tpx1^C48S^, even in the absence of cellular stress (**Fig. 3A, 3C**). Importantly, phosphatase-treatment confirmed that this change in electrophoretic mobility was due to phosphorylation (**Fig. 3C**). This suggests that Sty1-Tpx1 complex formation is sufficient to promote phosphorylation of Wis1, even in the absence of stress.

### Sty1-Tpx1 complexes support Wis1 activation in absence of Mcs4 scaffold protein

These data raised the intriguing possibility that Tpx1 might act as a scaffold to promote signaling between components of the Sty1 signaling pathway. Mcs4 is a fungal-specific protein that performs a scaffold function, guiding the interaction of Wak1 and Win1 MAP3Ks with Wis1 (Morigasaki and Shiozaki, 2013)(**Fig. 1A**). Consistent with this essential function in supporting Sty1 activation, Δ*mcs4* mutant cells are delayed in their entry to mitosis; with cells reaching a significantly longer size than wild-type cells before dividing (**Fig. 4A**) (Buck et al., 2001; Morigasaki and Shiozaki, 2013; Shiozaki *et al*., 1997). To test whether Tpx1 could also act as a signaling scaffold, we examined whether Sty1-Tpx1 complex formation could bypass the requirement for Mcs4 for MAP3K-dependent phosphorylation of Wis1. Remarkably, analysis of cells obtained from a cross between strains expressing *sty1-tpx1* or Δ*mcs4*, revealed that Sty1-Tpx1 expression stimulated similar levels of constitutive Wis1 phosphorylation in Δ*mcs4* mutant as in wild-type (*mcs4^+^*) cells (**Fig. 4B**). Accordingly, consistent with expression of the Sty1-Tpx1 fusion completely bypassing the requirement for Mcs4 for Sty1 activation, there were similar levels of Sty1-Tpx1 phosphorylation in *sty1-tpx1 mcs4*^+^ and *sty1-tpx1 Δmcs4* strains (**Fig. 4B**). Moreover, loss of *mcs4* had no effect on the length of septating cells expressing Sty1-Tpx1; both *sty1-tpx1 mcs4*^+^ and *sty1-tpx1 Δmcs4* cells were significantly shorter than wild type septating cells. These data demonstrate that constitutive Sty1-Tpx1 complex formation is sufficient to increase Wis1/Sty1 activity and accelerate entry into mitosis even in the absence of Mcs4 (**Fig. 4A-B**).

**Figure 4:**
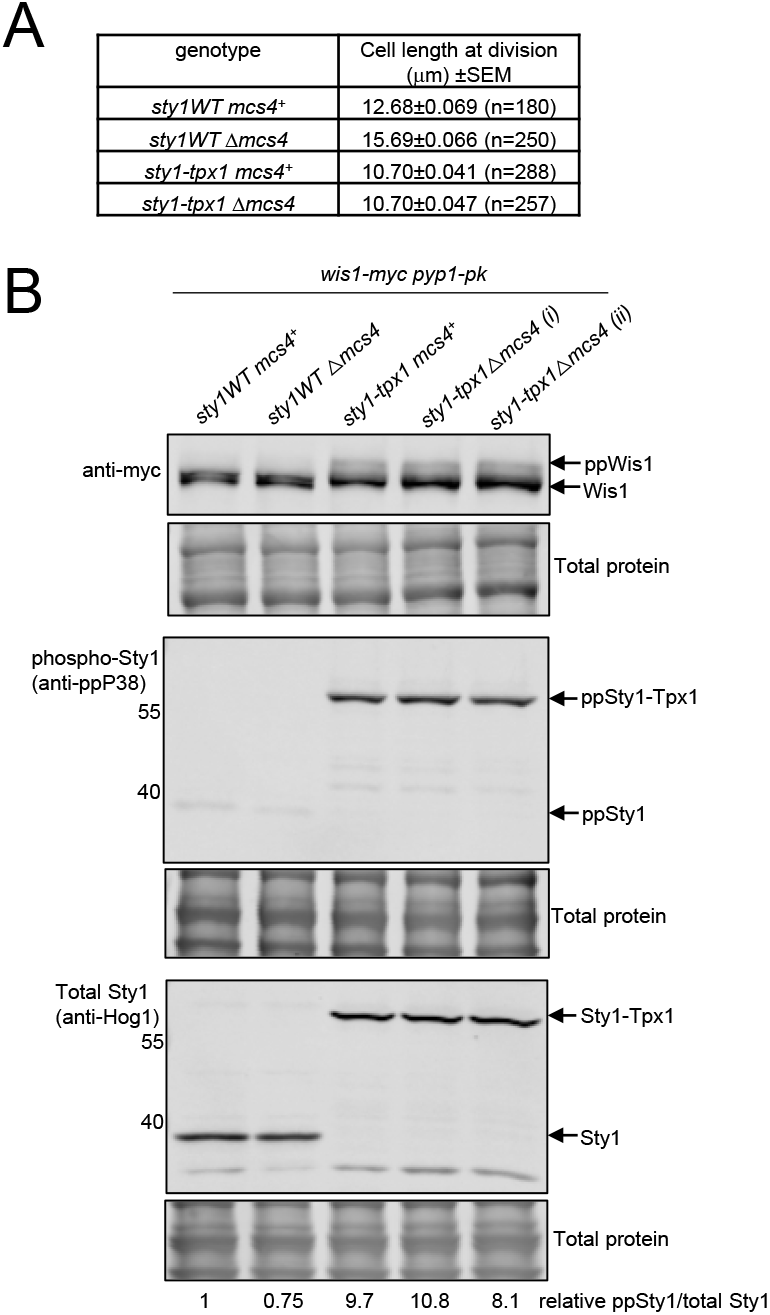
Sty1-Tpx1 complexes bypass the requirement for Mcs4 for phosphorylation of Wis1 and activation of Sty1. **[A]** Expression of Sty1-Tpx1 fusion protein rescues the long cell phenotype of Δ*mcs4* cells; Table shows the mean cell length of dividing cells ± Standard Error of the mean (SEM) determined by measuring the length of the indicated number of calcofluor-stained septating *WT* (MC2), Δ*mcs4* (MC77), *sty1-tpx1* (MC12), and *sty1-tpx1 Δmcs4* (MC83) cells. **[B]** Immunoblotting analysis of the phosphorylation and levels of Wis1 and Sty1 in cell lysates from *WT* (MC2), Δ*mcs4* (MC77), *sty1-tpx1* (MC12), and two different *sty1-tpx1 Δmcs4* (MC83) strains expressing Wis1-12myc. Blots were analyzed with anti-myc, anti-ppP38 and anti-Hog1 antibodies and total protein stain. Relative levels of phosphorylated Sty1, normalized to total Sty1, are shown below each lane.

### Tpx1-Sty1 complexes provide a protective scaffold for Wis1

The ability of a constitutively formed complex between Tpx1 and Sty1 to promote Wis1 phosphorylation, even in the absence of stress, suggests that H_2_O_2_-induced Tpx1-Sty1 disulfide formation activates Wis1 and Sty1 concomitantly. Thus we considered the possibility that the formation of disulfide complexes with Tpx1 might generate a ‘signaling island’, enhancing interactions between components of the Sty1 MAPK signaling pathway. Our analysis revealed that in cell extracts prepared under native conditions, Wis1 was sensitive to cleavage and partial degradation (**Fig. 5A**). Strikingly, although Sty1-Tpx1 and wild-type cells express similar total levels of Wis1 (**Figs. 3, 4, 5A**), much greater levels of ‘soluble’, intact Wis1 were present in native protein extracts prepared from Sty1-Tpx1 expressing cells (**Fig. 5**). This strongly suggests that Sty1-Tpx1 complexes protect Wis1 in native extracts from proteolytic degradation. Intriguingly, the low levels of Wis1 in cells expressing a Sty1^5CS^-Tpx1 mutant suggested that cysteines in Sty1 may be important for this protective function of Sty1-Tpx1 complexes (**Fig. 5A-C**).

**Figure 5:**
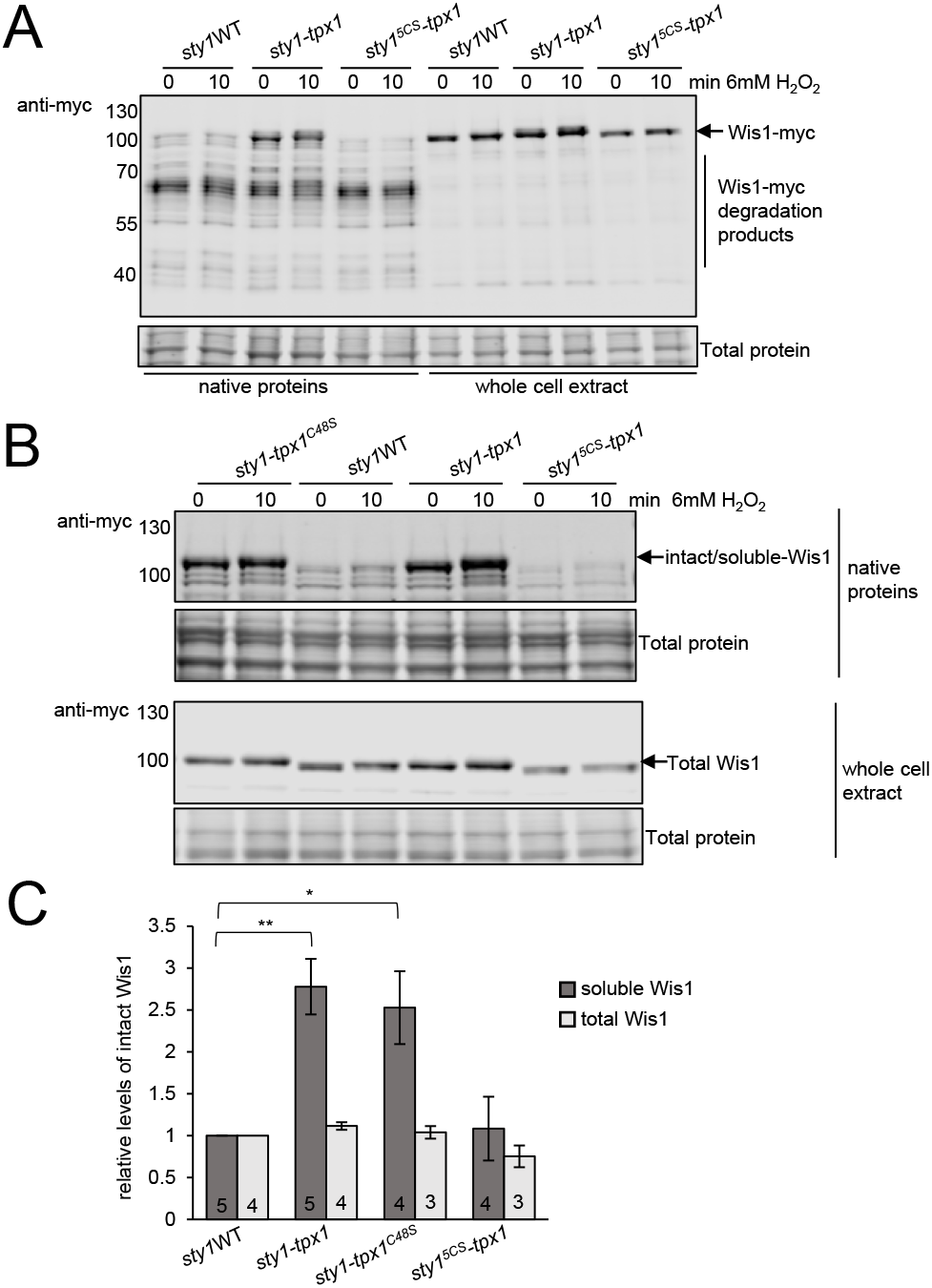
Sty1-Tpx1 complexes stabilize Wis1 and this is dependent on the presence of cysteines in Sty1. Immunoblotting analysis with anti-myc, anti-ppP38 or anti-Hog1 antibodies, of native (soluble proteins) and whole cell lysate prepared from cells expressing **[A]** wild-type sty1 WT (MC2), *sty1-tpx1* (MC12) and *sty^5CS^-tpx1* (MC13) strains or **[B]** wildtype Sty1 WT (MC2) *sty1-tpx1* (MC12), sty1-tpx1^*C48S*^ (MC82) and *sty^5CS^-tpx1* (MC13) before and after treatment for 10 min with 6 mM H_2_O_2_. **[C]** Bar chart quantification of data from multiple experiments indicates that although total Wis1 levels are similar, the levels of intact soluble Wis1 are 2.5x higher in cells expressing Sty1-Tpx1 (MC12) (t test p=0.0058) and Sty1-Tpx1^C48S^ (MC82) (t test: p=0.039) fusion proteins than wild-type Sty1 (MC2) or Sty1^5CS^-Tpx1 (MC13). Error bars indicate the mean level of protein ± Standard Error of the Mean (SEM) as determined from 3-5 repeated experiments (as indicated by numbers on bars in panel). See also Figure S5.

It was possible that increased Wis1 stability correlated with increased Wis1 phosphorylation. However, when cells were exposed to levels of H_2_O_2_ that induced Wis1 phosphorylation, we did not detect any consistent change in Wis1 stability (**Fig. 5A-B**). Moreover, analysis of Sty1-Tpx1^C48S^ expressing cells revealed that the peroxide-reacting cysteine of Tpx1 was dispensable for the stabilization of Wis1 (**Fig. 5B-C**). Together, these data suggested that Sty1-Tpx1 complexes may protect Wis1 from degradation by a mechanism that does not require Tpx1’s peroxidase activity and that this may be responsible the increased levels of Wis1/Sty1 activity observed in these cells (**Figs. 1, 3,4, 5)**.

### Wis1 is regulated by MAP3K-independent autophosphorylation

Next, we considered how Tpx1-Sty1 complexes might stabilize Wis1 and increase its phosphorylation. To further investigate how Wis1 phosphorylation is regulated, we made use of a chemical genetic approach using Δ*pyp1*Δ*pyp2* cells expressing Wis1DD and an ATP analogue-sensitive Sty1 mutant (Sty1^T97A^). The viability of these cells is maintained by inhibiting Sty1 kinase activity with the ATP analogue, 3-BrB-PP1, allowing activation of Sty1 in a stress-independent manner by removal of the drug (Mutavchiev et al., 2016). The small but reproducible stress-induced decrease in Wis1DD mobility we observed in these cells was consistent with our other data indicating that Wis1 undergoing a stress-induced phosphorylation on different sites from those phosphorylated by the MAP3K (**Fig. 6A, S4**). Importantly, this experiment also eliminated the possibility that this stress-induced phosphorylation of Wis1 involved inhibition of either Pyp1 or Pyp2, as neither phosphatase was present in these cells (**Fig. 6A**). Furthermore, the inhibition of Sty1’s kinase activity by 3-BrB-PP1, also eliminated the possibility that Sty1’s kinase activity was responsible for these stress-induced increases in Wis1 phosphorylation (**Fig. 6A, S5A-B**). Similarly, analysis of cells expressing an analogue-sensitive version of Sty1 fused to Tpx1 (Sty1^T97A^-Tpx1) suggested that inhibiting the kinase activity of the Sty1^T97A^-Tpx1 fusion (with 3-BrB-PP1) did not reduce the level or extent of Wis1 phosphorylation (**Fig. S5C**). Based on these data we concluded that the stress-induced, Tpx1-dependent phosphorylation of Wis1, that is increased by Sty1-Tpx1 complexes, does not require Sty1’s kinase activity or the MAPK phosphatases Pyp1 and Pyp2.

**Figure 6:**
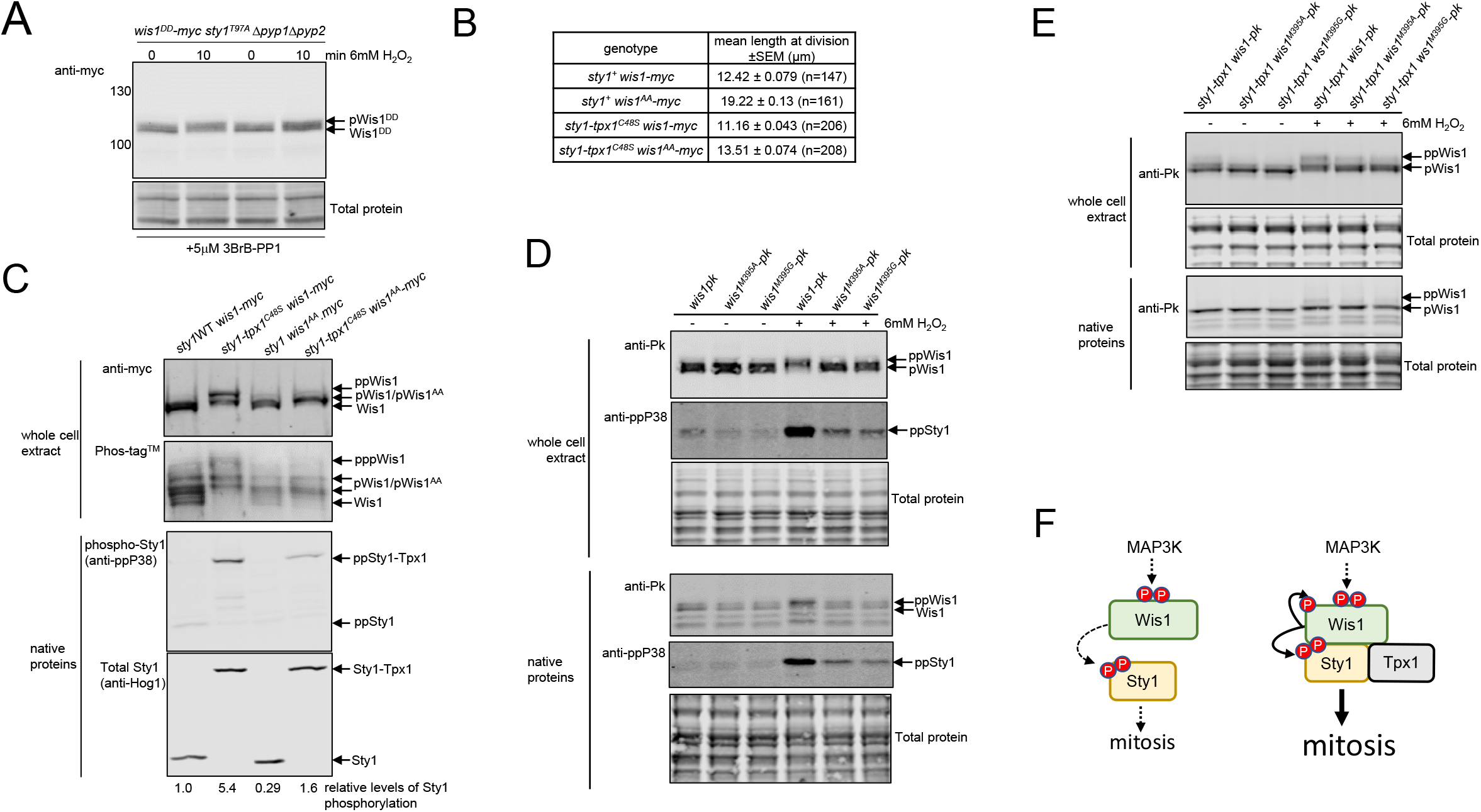
Wis1 undergoes autophosphorylation in response to stress or in presence of Sty1-Tpx1 complexes [A]. Immunoblot analysis, using anti-myc antibodies of the mobility of Wis1^DD^.myc in Δ*pyp1* Δ*pyp2 Sty1*^T97A^ (SISA) cells (lanes 1 and 2: KS8266 and lanes 3 and 4: KS8311) (Mustavchiev et al., 2016) grown in EMM media with supplements containing 5μM 3-BrB-PP1, washed 2x in either EMM or EMM containing 5μM 3BrB-PP1, then transfered to fresh EMM or EMM containing 5μM 3-BrB-PP1, before treatment (where indicated) with 6mM H_2_O_2_. **[B-C]** Expression of a Sty1-Tpx1 fusion is sufficient to rescue phenotypes associated with loss of MAP3K-dependent phosphorylation of Wis1 **[B]** Table shows the mean length of cells expressing wild-type Sty1 and Wis1 (MC132), wild-type Sty1 or Sty1-Tpx1^C48S^ fusion and wild-type Wis1 or Wis1^AA^: *sty1*^+^ *wis1^AA^-myc* (MC122), *sty1-tpx1^C48S^* (MC135) and *sty1-tpx1^C48S^ wis1^AA^-myc* (MC130). **[C]** Immunoblotting analysis with anti-myc, anti-ppP38 or anti-Hog1 antibodies of native proteins and whole cell lysate prepared from cells expressing wild-type Wis1.myc and Sty1 (MC132) or cells expressing *wis1^AA^-myc* (MC122), *Sty1-Tpx1^C48S^ and Wis1-myc* (MC135) or *Sty1-Tpx1^C48S^ and wis1^AA^-myc*, separated, as indicated, by Phos-tag^TM^ or conventional SDS-PAGE. **[D-E]** Immunoblotting analysis with anti-Pk, anti-ppP38 or anti-Hog1 antibodies of native and whole cell lysate prepared from cells expressing wild-type Wis1-3pk, Wis1^M395A^-3pk or Wis1^M395G^-3pk with **[D]** wild-type Sty1 (MG46, MG47, MG48) or **[E]** Sty1-Tpx1 fusion protein (MG49, MG50, MG51), before or following treatment for 10 min with 6 mM H_2_O_2_ **[F]** Model illustrating how Tpx1 provides a scaffold/signaling platform supporting the stress-induced activation of Wis1 by MAP3K and autophosphorylation. Thus, constitutive ‘prdxylation’ of the Sty1 MAPK accelerates entry into mitosis. See also Figure S6.

To further investigate this phosphorylation of Wis1, we expressed Sty1-Tpx1^C48S^ fusion in cells expressing a Wis1^AA^ mutant, in which the MAP3K-phosphorylated residues are substituted with alanine. Wis1^AA^ cells are significantly elongated reflecting the mitotic delay associated with severely reduced levels of Sty1 phosphorylation (Shiozaki et al., 1998) (**Fig. 6B, S6A**). Significantly, consistent with Sty1-Tpx1 acting to increase Wis1 activity independently from MAP3K-dependent phosphorylation, expression of Sty1-Tpx1^C48S^ increased Sty1 phosphorylation and partially rescued the cell cycle defect of Wis1^AA^ cells (**Fig. 6B-C**). However, a smaller increase in Wis1^AA^ phosphorylation was observed in cells co-expressing Sty1-Tpx1^C48S^ compared with the increased phosphorylation of wild-type Wis1 (compare lanes 2 and 4 in **Fig. 6C**). As Wis1^AA^ has reduced kinase activity compared with wild-type Wis1, this raised the possibility that Wis1 kinase activity might be important for the non-canonical phosphorylation detected following stress or Sty1-Tpx1 complex formation. To examine this possibility, we generated cells expressing ‘gatekeeper’ mutant versions of Wis1, Wis1^M395G^ or Wis1^M395A^, which are predicted to be sensitive to inhibition by the ATP analogue 3-BrB-PP1 (Gregan et al., 2007). Although, both Wis1^M395G^ and Wis1^M395A^ were expressed at wild-type levels, Sty1 phosphorylation was very low and minimally increased in response to stress in cells expressing either Wis1^M395G^ or Wis1^M395A^ (**Fig. 6D**). This suggested that the catalytic activity of both Wis1 mutant proteins was severely reduced, even in the absence of the ATP analogue. Moreover, consistent with the requirement of Wis1-dependent phosphorylation of Sty1 for timely entry into mitosis, cells expressing Wis1^M395G^ or Wis1^M395A^ were longer than cells expressing wild-type Wis1 (**Fig. S6B**). Strikingly, the stress-induced phosphorylation of Wis1^M395G^ and Wis1^M395A^ was also compromised compared with wild-type Wis1 (**Fig. 6D**). This strongly suggests that Wis1 kinase activity is required for the stress-induced phosphorylation and activation of Wis1. Similarly, the phosphorylation of Wis1 in cells expressing Sty1-Tpx1 fusion proteins was less extensive in cells expressing either Wis1^M395G^ or Wis1^M395A^ **(Fig. 6E**). These data suggest that in response to stress, Sty1-Tpx1 complex formation promotes the autophosphorylation of Wis1 and that this contributes to the increased activity of Wis1 towards Sty1 under these conditions. Based on these data we propose that Tpx1-Sty1 (Prdx-P38) complexes behave as a molecular scaffold, promoting activation of the MAPKK Wis1 and entry into mitosis (**Fig. 6F**).

In summary, our data suggest that the Prdx, Tpx1, facilitates H_2_O_2_-dependent activation of Sty1 (p38) by two mechanisms (1) by promoting thioredoxin-dependent oxidation of the MAPK tyrosine phosphatase, Pyp1, and (2) by forming disulfide complexes with the Sty1 MAPK (**Fig. 7A**). We show that Tpx1-Sty1 (Prdx-P38) complexes provide a scaffold supporting stable MAPKK-MAPK interactions and the canonical and non-canonical phosphorylation of the MAPKK, Wis1 (**Fig. 7**). As depicted in our model (**Fig. 7B**), we propose these mechanisms confer H_2_O_2_-sensitivity on the P38 (Sty1) MAPK; facilitating the activation of P38 (Sty1) by H_2_O_2_ levels far below the threshold for activating the canonical MAP3K.

**Figure 7.**
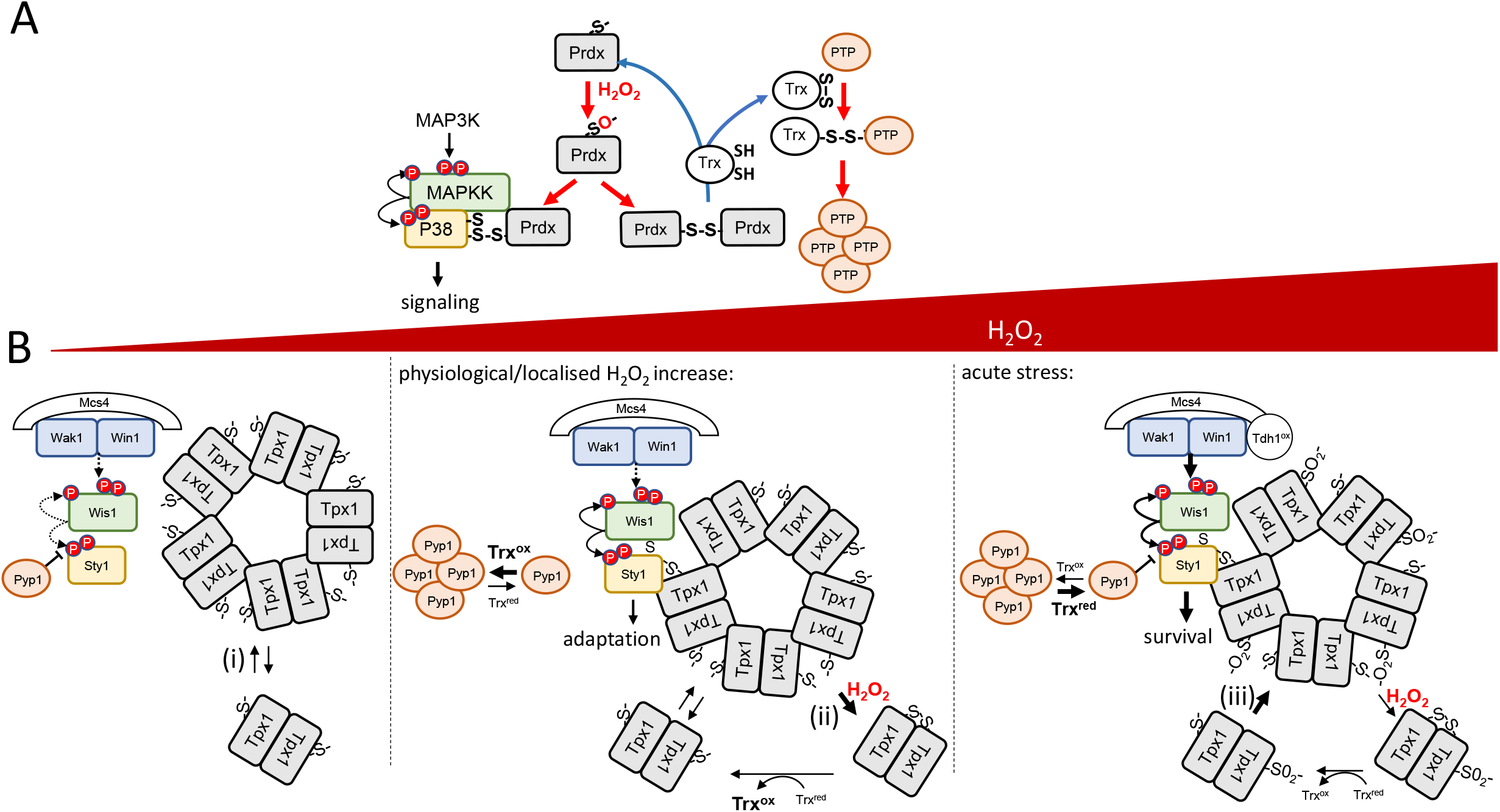
Models illustrating how Prdx-disulfide complexes can act as mediators of H_2_O_2_ signal transduction to the P38/Sty1 pathway [A]. We propose that Prdx (Tpx1) acts as a peroxide transducer to increase activation of P38/Sty1 pathway by two independent mechanisms. (1) Prdx-Sty1 MAPK complexes stabilize interactions with the MAPKK Wis1 to provide a MAPKK-MAPK signaling island that allows increased autophosphorylation and canonical phosphorylation of the MAPKK, Wis1. (2) The reduction of Prdx-Prdx disulfides promotes oxidation of thioredoxin (Trx) (Day et al 2012) and thus promotes the formation of disulfide complex by the MAPK phosphatase (PTP) Pyp1. **[B]** We propose that the H_2_O_2_-induced formation of complexes between the Prdx, Tpx1, and P38/Sty1, provides a signaling scaffold, that supports the increased activation of Wis1 by non-canonical autophosphorylation and MAP3K-dependent canonical phosphorylation to ensure the appropriate response to different levels of H_2_O_2_: We propose that Prdx oligomers may contribute to this scaffold as described below: (i) Prdx dimers like Tpx1 assemble into decamers (For a review see Bolduc et al., 2021). (ii) H_2_O_2_-induced formation of Tpx1-Tpx1 disulfide bonds at low levels of H_2_O_2_ is predicted to promote dissociation of Tpx1 decamers into dimers. Therefore, under these conditions we propose that Tpx1-dependent, thioredoxin-mediated oxidation of the Pyp1 phosphatase into higher order disulfide-linked complexes, may be important to inhibit Pyp1 and increase Sty1 activity. (iii) As the cell’s H_2_O_2_-buffering capacity becomes saturated (Tomalin et al., 2016) the peroxidatic cysteine of Tpx1 becomes hyperoxidized to sulfinic (SO_2_^-^). This is predicted to stabilize non-covalent interactions between Tpx1 dimers (Noichri et al., 2015). Accordingly, we predict this could increase the stability of the Tpx1 signaling scaffold, increasing Wis1 stability, autophosphorylation and canonical phosphorylation by the MAP3K (Wak1 and Win1). At high concentrations of H_2_O_2_, oxidation of the GAPDH (Tdh1) increases Sty1 activation by increasing MAP3K activity (Morigasaki et al., 2008). We propose that the increased availability of reduced thioredoxin following hyperoxidation of Tpx1 at higher concentrations of H_2_O_2_ (Day et al., 2012) may contribute to negative feedback mechanisms to restore Pyp1 activity.

## Discussion

Here we demonstrate that formation of a complex with a Prdx can be sufficient to increase the activity of P38 MAPK. Our data suggests the Prdx-MAPK complex provides a scaffold supporting increased MAP3K-MAPKK-MAPK signal transduction, particularly the increased activation of the MAPKK by canonical and non-canonical phosphorylation (**Fig. 6F, 7**). Intriguingly, our genetic fusion experiments suggest that the Prdx-P38 scaffold does not require the peroxide-reacting cysteine of the Prdx and operates at H_2_O_2_ concentrations above those at which this cysteine is hyperoxidized to sulfinic (SO_2_^-^) (Veal et al., 2004; Bozonet et al., 2005; Vivancos et al., 2005; Day et al., 2012). Moreover, our discovery that fusion to a Prdx increases the phosphorylation of human P38 (**Fig. 1H**) suggests this peroxidase-independent scaffold function for Prdx in activating P38 MAPK is conserved. However, Prdx dimers readily assemble into decamers (For a review see Bolduc et al., 2021) and it is possible that oligomerization with endogenous Prdx may be important for Prdx-P38 scaffold function (**Fig. 7B**). In which case, it is possible that Prdx hyperoxidation (SO_2_^-^), which promotes Prdx oligomerization (Jang et al., 2004; Noichri et al., 2015; Bolduc et al., 2021), could stabilize the Prdx-P38 scaffold, potentially increasing the duration of P38 activation at higher concentrations of H_2_O_2_ (**Fig. 7B**).

Prdx have been shown to form intermolecular disulfide complexes with several components of P38 MAPK signaling pathways (Barata and Dick, 2020; Jarvis *et al*., 2012; Veal *et al*., 2004). Based on the archetypal example of Prdx2-STAT3 disulfide complexes, which lead to further disulfides that inactivate STAT3 (Sobotta *et al*., 2015), these complexes have been proposed to function as intermediates; leading to the formation of additional disulfide bonds that alter activity. Here we have explored the alternative hypothesis that these Prdx-MAPK complexes themselves have altered function (van Dam et al., 2021). Our molecular genetic approach, with cells engineered to express P38-Prdx fusions, has established that complexes between Prdx and P38 MAPK alter P38 activity by providing a platform for increased MAPKK-MAPK signaling. This is reminiscent of the role that covalent post-translational modification of proteins, for example, by the Small Ubiquitin-Like Modifier, SUMO, can play in regulating protein activity and signal transduction (Pelisch et al., 2017; Verhelst et al., 2011). Notably, many proteins have been identified to form H_2_O_2_-induced disulfide complexes with Prdx (Stocker et al., 2018; van Dam *et al*., 2021). It remains to be determined whether our discovery that the formation of complexes with a Prdx is sufficient to alter the activity of P38 signaling proteins is replicated for other proteins that form disulfide-bonded complexes with Prdx. However, if this is the case, then perhaps the formation of H_2_O_2_-induced disulfide complexes with Prdx should be considered as a redox-regulated post-translational modification. Significantly, Prdx can also act as ATP-independent molecular chaperones, protecting cells against protein aggregation under acute stress conditions (For a review see Ulrich et al., 2021). Thus, it is tempting to speculate that the formation of disulfide complexes with Prdx may identify specific targets for this chaperone activity. Indeed, our finding that interactions between a MAPK and a Prdx increase the stability of the associated MAPKK is notable in this context; raising the possibility that disulfide complexes between Prdxs and signaling proteins may also increase the stability of other complexes, or components of the complexes, in which they participate. Intriguingly, the inability of Sty1^5CS^-Tpx1 fusion to increase Wis1 levels or phosphorylation suggests that cysteines in Sty1 are also needed for the MAPKK-MAPK signaling platform. The conservation of these cysteines in other MAPK, raises the possibility that redox-active surface cysteines may be important for maintaining interactions in other MAPKK-MAPK signaling modules.

Our genetic evidence has indicated a second, independent mechanism through which the peroxiredoxin Tpx1 can promote the H_2_O_2_-induced activation of the P38-related Sty1 MAPK: We find that high levels of Tpx1 promote the activation of Sty1 in response to low levels of H_2_O_2_, by promoting the transient, thioredoxin-dependent, oxidation of the MAPK phosphatase, Pyp1, into high molecular weight disulfide complexes. The molecular nature of these complexes remains unclear, and it is possible that they could contribute to the signaling scaffold. However, our data are also consistent with these complexes lacking Pyp1 phosphatase activity. Protein tyrosine phosphatases (PTPs) are well-established to be regulated by reversible oxidation of the invariant catalytic cysteine, with formation of intramolecular disulfide bonds representing an established mechanism for preventing irreversible oxidation of the catalytic cysteine to sulfinic or sulfonic forms (Tonks, 2005).

Pyp1 has previously been shown to be susceptible to inhibition by stress-induced aggregation, providing a mechanism for MAP3K-independent activation of Sty1 in response to heat stress (Nguyen and Shiozaki, 1999; Samejima et al., 1997; Boronat *et al*., 2020). Redox-regulated protein-protein interactions can also influence the duration of PTP inhibition (Londhe et al., 2020; Turner-Ivey et al., 2013). Thus we propose that the Prdx/Trx-dependent disulfide complexes Pyp1 forms in response to H_2_O_2_ may be important to allow transient, reversible inactivation of Pyp1 (**Fig. 7**).

As discussed above, Prdx have been shown to directly participate in the H_2_O_2_-induced oxidation of cysteines in some signaling proteins. The oxidoreductase activity of the thioredoxin, Trx1, is important for reducing disulfide bonds in many proteins, including Tpx1 (Day *et al*., 2012). Hence, if Tpx1 participated directly in the formation of Pyp1 disulfide complexes, one might have expected that loss of Trx1 would *increase* Pyp1 oxidation. In contrast, we have found that formation of these complexes absolutely requires Trx1, which rapidly (within 30s) forms a H_2_O_2_-induced disulfide complex with Pyp1. Moreover, we could find no evidence for disulfide complexes between Tpx1 and Pyp1. These data suggest that Pyp1 oxidation does not directly involve Tpx1 as a peroxide transducer. Instead, they suggest that high levels of Tpx1 promote Pyp1 oxidation to disulfide-containing complexes indirectly, by promoting the oxidation of Trx1 (Day *et al*., 2012). The thioredoxin peroxidase activity of Tpx1 has been shown to promote Pap1 oxidation by promoting Trx1 oxidation (Day *et al*., 2012). This was previously suggested to reflect the need to inhibit thioredoxin-mediated reduction of Pap1. However, we note that a direct, pro-oxidant function for oxidized Trx1 would be reminiscent of the role of thioredoxin-family proteins, such as DsbA or protein disulfide isomerase (PDI), in initiating disulfide bond formation during the oxidative folding of periplasm/ER/secreted proteins: These initial steps in oxidative protein folding involve oxidized DsbA, in prokaryotes, or thioredoxin domains in PDI, accepting electrons from protein-thiols to form intermolecular disulfide bond intermediates (Bulleid and Ellgaard, 2011). It is tempting to speculate that the Pyp1-Trx1 disulfides we observe play a similar role in promoting additional Pyp1-disulfide formation/oxidation events. Notably, the critical role of Tpx1 in the formation of Pyp1-disulfide complexes is also reminiscent of the role that the ER-based Prdx, Prdx4, can play in promoting oxidative protein folding; where Prdx4 disulfides have been shown to act as an electron acceptor for PDI in mammalian cells lacking Ero1 (Bulleid and Ellgaard, 2011; Tavender et al., 2010; Zito et al., 2010). Similarly, in cells exposed to an H_2_O_2_ bolus Tpx1 disulfides are highly abundant, rendering them an electron acceptor that leads to thioredoxin oxidation (Day *et al*., 2012). There is a growing appreciation that localized increases in endogenous H_2_O_2_ can initiate physiological responses. Prdx have been shown to be directly involved in some of these responses (For reviews see Winterbourn, 2020; Bolduc et al., 2021). Although, here we have applied an external bolus of H_2_O_2_, our data suggest that Prdx-driven localized oxidation of thioredoxin could also provide a means to facilitate relay of endogenous H_2_O_2_ signals to nearby signaling proteins.

The P38-related MAPK Sty1, like its counterparts in other eukaryotes responds to a multitude of extracellular stimuli. In *S. pombe* there are only two MAP3K, Wak1 and Win1, and a single MAPKK, Wis1, that is required for Sty1 phosphorylation and activation. However, the mechanisms by which these kinases are activated in response to particular metabolic and stress stimuli are not well established. One aspect of Sty1 activation involves the Mcs4 protein, which acts as an essential scaffold; supporting MAP3K activity and Sty1 activation by stabilising interactions of the MAP3K, Wak1 and Win1 (Morigasaki and Shiozaki, 2013). The glyceraldehyde phosphate dehydrogenase (GAPDH) Tdh1, and two component signaling proteins regulating Mcs4, have also been shown to be important for promoting MAP3K activity in response to H_2_O_2_ (Buck *et al*., 2001; Morigasaki et al., 2008; Shiozaki *et al*., 1997). However, we have previously shown that Tpx1 acts downstream of Mcs4 to promote H_2_O_2_-induced activation of Sty1 (Veal *et al*., 2004). We now demonstrate clearly that Sty1-Tpx1 complexes are able to substitute for Mcs4’s scaffold function, restoring Sty1 activity to *□mcs4* mutant cells. Moreover, Sty1-Tpx1 complexes also increase Sty1 activity in cells where MAP3K-dependent activation is prevented by mutation of the canonical MAP3K sites in the MAPKK Wis1. Consistently, MAP3K-dependent increases in Wis1 phosphorylation do not occur until much higher concentrations of H_2_O_2_ than those required for Pyp1-Trx1 and Sty1-Tpx1 disulfide formation, or Sty1 activation. This indicates the key role that Tpx1-dependent oxidation events play in facilitating the important Sty1-mediated adaptive responses to lower levels of H_2_O_2_. Moreover, we have demonstrated that Tpx1-driven oxidation of the protein tyrosine phosphatase Pyp1 and Sty1-Tpx1 complexes act by independent mechanisms, synergistically activating Sty1. This combination of Tpx1-dependent mechanisms to simultaneously promote MAPKK activation and inhibit the MAPK phosphatase Pyp1, provides an attractive mechanism to tailor a rapid response to small increases in H_2_O_2_ at which MAP3K are not activated. Intriguingly, it has also been suggested that the upstream activators of P38 in mammalian cells, the MAPKK MKK3 and MKK6, can be modulated through intramolecular or intermolecular disulfide signaling to P38 suggesting aspects of this regulation may be conserved (Bassi et al., 2017; Diao et al., 2010).

We also reveal that Wis1 undergoes an additional stress-induced, non-canonical phosphorylation that is stimulated by the presence of Sty1-Tpx1 complexes. Although we have not determined the precise nature of this phosphorylation event, our analysis of cells expressing Wis1 mutants with greatly reduced kinase activity are consistent with it reflecting autophosphorylation by Wis1. It will be interesting to determine how this autophosphorylation contributes to the increased stability and activity of Wis1 in cells expressing Sty1-Tpx1 fusion proteins. Given the intense interest in therapeutic targeting of the P38 pathway, this discovery could also suggest new avenues for modulation of this signaling node. A major challenge for the clinical use of P38 inhibitors, is specifically targeting cells that are dependent on elevated P38 activity, avoiding the unwanted side effects of broadly inhibiting P38 activity in all cells (Martinez-Limon et al., 2020). Here we demonstrate that constitutive P38-Prx complex formation represents a non-canonical mechanism to activate P38 MAPKs, which is lethal when combined with loss of MAPK phosphatase activity. This could invite new strategies to specifically target P38 signaling outputs in tumor cells with elevated Prdx levels, or in which P38 is hyperactivated by the absence of normal feedback mechanisms.

## Acknowledgements

We would like to thank Professors Janni Petersen, Janet Quinn, Ken Sawin, Per Sunnerhagen and Dr Johanna Sjölander for kindly providing us with the indicated *S. pombe* strains. We are grateful to Professor Brian Morgan, Dr Josana Rodriguez and Dr Paraskevi Kritsiligkou for providing valuable comments on the manuscript. We thank the BBSRC for funding BB/T002484/1 (MC, EB and EV), BB/F023065/1 (AD, EV), BBSRC DTP studentships (BB/M011186/1 to MG and EV.), (BB/J014516/1 to HL and EV), BB/T008695/1 (to ED and EV). P.A.E. and D.P.B. acknowledge funding from the NIH Illuminating the Druggable Genome (IDG) common fund consortium (NCI U01CA239106), BBSRC grants BB/S018514/1 and BB/N021703/1 and North West Cancer Research (NWCR) grant CR1208.

## Author Contributions

Conceptualization, EV, MG; Methodology, EV, MG, MC, DB, PE, HL, ED; Validation, MC, HL, MG, DB; Investigation, MC, AD, MG, HL, DB, EB; Resources, ZU; Writing -Original draft, EV, MC, MG, AD, DB, PE; Writing -Review & Editing, EV, MC, MG, AD, DB, PE; Supervision, EV and PE; Project Administration, EV; Funding Acquisition, EV and PE.

## Experimental Procedures

### STAR METHODS

#### RESOURCE AVAILABILITY

Lead contact Further information and requests for resources and reagents should be directed to and will be fulfilled by the lead contact, Elizabeth Veal

##### Materials availability

Yeast strains (Table 1) and plasmids (Table 2) generated by this study are available on request from the lead author.

##### Data and code availability

All data in this paper will be shared by the lead contact upon request.

This paper does not contain code.

Any additional information required to re-analyze data will be supplied by the lead contact upon request.

#### EXPERIMENTAL MODEL AND SUBJECT DETAILS

##### Yeast strains, growth conditions and transformation

The *S. pombe* strains used in this study are shown in **Table 1**. Cells were maintained on Ye5S or Edinburgh minimal medium 2(EMM2) including appropriate supplements (Moreno et al., 1991). SISA strains were maintained on Ye5S media containing 5μM 3-BrB-PP1(Abcam, ab143756) to prevent lethal activation of sty1 and sequenced to confirm continued presence of Wis1^DD^-expressing allele (Mustavchiev et al., 2016). For experiments, cells were grown in liquid culture at 30 °C to mid-log phase (OD_595_=0.3-0.5) with constant aeration at 180rpm in Edinburgh minimal medium 2(EMM2) with appropriate supplements, unless otherwise indicated. DNA was introduced into cells by lithium acetate based chemical transformation.

##### Mammalian cell growth and transfection

HEK-293T cells were cultured in Dulbecco’s modified Eagle medium (Lonza) supplemented with 10% fetal bovine serum (HyClone), penicillin (50 U/ml), and streptomycin (0.25 μg/ml) (Lonza) and maintained at 37°C in 5% CO_2_ humidified atmosphere. HEK-293T cells were transfected using a 3:1 polyethylenimine (PEI [branched average *M*_w_ ~25,000 Da; Sigma-Aldrich]) to DNA ratio (30:10 μg, for a single 10-cm culture dish).

#### METHOD DETAILS

##### Spot tests

An equal number of exponentially growing cells were serially diluted 10-fold and spotted on to Ye5S containing 0.4 mM H_2_O_2_, 1 mM H_2_O_2_ or 0.5 M KCl. Plates were incubated at 30°C for 2-4 days before imaging.

##### Plasmid and strain construction

*Construction of Sty1-Tpx1 expressing gene fusion strains expressed from S. pombe sty1 locus*. pRip2*tpx1+* and *pRip2tpx1^C48S^* were generated by removing ARS region from pRep2*tpx1* or *pRep2tpx1^C48S^* using *EcoRI* (Veal et al., 2004). The *sty1^+^, sty1^5CS+^* and *sty1^T97A^* genes including 1.5kb of the *sty1* promoter up stream of the open reading frames were amplified by Phusion (Thermo Scientific) PCR from NT5, AD84 and KS7830 genomic DNA respectively using primers containing *PstI* and *NdeI* restriction sites at 5’ end and 3’ end respectively (**Table 2**). The amplified PCR products were cloned into *PstI* and *NdeI* sites of pRip2*tpx1*^+^ or pRip2*tpx1*^C48S^. The recombinant plasmids were linearised with *NheI* and transformed in to AD22. The sequences of sty1 promoter, *sty1/sty1^5CS^/sty1^T97A^* ORF were confirmed by sequencing (Eurofins) of DNA from URA+ transformants using primers *PstI* sty1 mutant N and sty1 internal F.

*Construction of S. pombe expressing Pk-tagged Pyp1 from the pyp1 locus* The *pyp1*^+^ orf *was amplified* by PCR using primers containing Pst1 and BamH1 sites and ligated into Pst1/BamHI digested pRip42PkC to create pRip42Pyp2PkC. This was linearised with SaI1 and transformed into wild-type (NT4) cells. URA+ colonies were screened for integration at *pyp1*^+^ locus by colony pcr and sequenced with Pyp1intcheck and Pyp1intcheckB primers.

*Construction of S. pombe expressing Pk-tagged Wis1, wis1^M395A^ and wis1^M395G^ mutant proteins from the wis1 locus* Mutations of gate-keeper residue Methionine 395 in wis1 ATP-binding pocket were designed to confer sensitivity to small-molecule inhibitors (Gregan et al., 2007). Wild type *wis1*^+^ open reading frame or versions in which Methionine at 395 was substituted with alanine or glycine were synthesised by integrated DNA technologies (IDT) with *PstI* site at 5’end and *KpnI* site at 3’end and cloned into pJet1.2 (Thermo Scientific). Wild type *wis1*^+^ ORF was amplified by Phusion PCR from NT4 genomic DNA with primers containing *PstI* and *KpnI* restriction sites at 5’ end and 3’ end respectively. Amplified PCR products were digested then ligated into *PstI* and *KpnI* sites of pRip42pk (Day et al., 2012) to generate pRip42Wis1pkC, pRip42Wis1^M395G^pkC or pRip42Wis1^M395A^pkC. Recombinant plasmids were linearised with *ApaI* and transformed into NT4 strain and URA+ colonies screened by colony pcr for integration at *wis1*^+^ locus with wis1_auto_*Pst/*_F and pk_tag_check_rev primers, then sequenced with wis1_mt395_chk_fwd primer.

###### Construction of P38-PRDX1/2 fusion plasmids for expression in mammalian cells

Synthetic genes for P38α/MAPK14 fused at the C-terminus to human PRDX1 or PRDX2 were custom synthesized by integrated DNA technologies (IDT) and provided in the pUCIDT-Kan GoldenGate vector. Constructs were designed to include a 5’ Kozak sequence consensus sequence, an N-terminal Flag-tag with a 3C protease cleavage site (LEVFLQG/P), a short linker region (GGGSGGG) between the two fusion proteins and a 3’ stop codon. P38α-PRDX1/2 genes were subcloned into the mammalian expression vector, pcDNA3 using BamHI and NotI (NEB) restriction enzyme and T4-ligase (NEB) following standard ligation protocols, and verified by sequencing (Eurofins). N-terminal Flag-tagged P38α containing 3C protease site was amplified by PCR using CloneAmp HiFi PCR Premix (Takara) and ligated in to pcDNA3 using the aforementioned restrictions sites. Cysteine to serine Prdx mutants were generated using standard PCR-based mutagenic procedures.

##### Analysis of Mammalian cellular proteins

Whole cell lysates were collected 48 h post transfection in bromophenol blue–free SDS-PAGE sample buffer supplemented with 1% (v/v) Triton X-100, protease inhibitor cocktail tablet, and a phosphatase inhibitor tablet (Roche), and sonicated briefly. Total cell lysates were clarified by centrifugation at 20,817 x *g* for 20 min at 4°C, and supernatants were sampled and diluted 30-fold to calculate protein concentration using the Coomassie Plus Staining Reagent (Bradford) Assay Kit (Thermo Fisher Scientific). Cell lysates were then normalised and processed for immunoblotting.

For immunoprecipitation experiments, proteins were harvested 48 h post transfection in a lysis buffer containing 50 mM Tris-HCl (pH 7.4), 150 mM NaCl, 0.1% (v/v) Triton X-100, 1 mM DTT, 0.1 mM ethylenediaminetetraacetic acid (EDTA), 0.1 mM ethylene glycol-bis(ß-aminoethyl ether)-*N*,*N*,*N*’, *N*’-tetraacetic acid (EGTA) and 5% (v/v) glycerol and supplemented with a protease inhibitor cocktail tablet and a phosphatase inhibitor tablet (Roche). Lysates were briefly sonicated on ice and clarified by centrifuged at 20,817*g* for 20 min at 4°C, and the resulting supernatants were incubated with anti-FLAG G1 Affinity Resin (GeneScript) for 1-3 hours (as required) with gentle agitation at 4°C. Affinity beads containing bound protein were collected and washed three times in 50 mM Tris-HCl (pH 7.4) and 500 mM NaCl and then equilibrated in storage buffer (50 mM Tris-HCl [pH 7.4], 100 mM NaCl, 1 mM DTT, and 5% (v/v) glycerol). The purified proteins were then proteolytically eluted from the suspended beads over a 1-hour period using 3C protease (0.5 μg) at 4 °C with gentle agitation. Purified protein was detected by western blotting using antibodies with specificity towards P38.

##### *In vitro p*rotein kinase assays using immunoprecipitated P38 or P38-Prdx fusion proteins

Nonradioactive kinase assays were performed using real-time mobility shift-based microfluidic assays, as described previously (Byrne *et al*., 2020), in the presence of 2 μM of the appropriate fluorescent-tagged peptide substrate (5-FAM-IPTSPITTTYFFFKKK-COOH), 1 mM ATP and 1 mM DTT. Pressure and voltage settings were adjusted manually to afford optimal separation of phosphorylated and nonphosphorylated peptides. All assays were performed in 50 mM Hepes (pH 7.4), 0.015% (v/v) Brij-35, and 5 mM MgCl_2_, and real-time peptide phosphorylation was calculated from the ratio of the phosphopeptide:peptide. Where specified, assays also included 10 μM of the selective P38 inhibitor SB239063 (Tocris)

###### Analysis of *S. pombe* proteins by immunoblotting

For native cell extracts: Equal numbers of approximating 2-2.5×10^8^ exponentially growing *S. pombe* were mixed with an equal volume of ice pelleted by centrifugation for 1 min then snap frozen in liquid nitrogen. Cell lysates were extracted in buffers containing protease and phosphatase inhibitors as previously described (Veal et al., 2002).

For whole cell extracts: Equal numbers of exponentially growing *S. pombe* (4.5×10^7^-5×10^7^) were added to 20% trichloroacetic acid (TCA), harvested by centrifugation and snap frozen in liquid nitrogen. Protein was extracted essentially as described previously (Delaunay et al., 2000) with precipitated proteins resuspended in TCA buffer (1% (w/v) SDS, 1 mM EDTA, 100 mM Tris-HCL pH 8.0). In experiments where oxidation state was examined e.g. Pyp1 in Fig. 2 proteins were dissolved in TCA buffer containing 10 mM *N*-Ethylmaleiminde (NEM) or, where indicated, 4’-acetamido-4’-maleimidylstilbene-2,2’-disulfonic acid (AMS) to prevent disulfide exchange and visualize any shifts due to cysteine oxidation. EDTA-free TCA buffer was used where phosphorylation of proteins was to be investigated by alkaline phosphatase-treatment (Roche) or Phos-tag^TM^ (NARD Institute, Japan) gels. Protein concentrations were estimated using the bicinchoninic acid protein assay (Thermo scientific) and equal amounts of denatured protein separated by SDS-PAGE on 8% acrylamide gels containing 100mM Phos-tag^TM^ where indicated. Separated protein was then transferred onto nitrocellulose membrane (GE healthcare) or, for Phos-tag gels, methanol-activated PVDF membrane (Immobilon-P Milipore). Where indicated, membranes were staining with Revert ^TM^ 700 total protein stain (Licor) for normalization of protein loading. Membranes were then blocked in TBS-T (15 mM NaCl, 1 mM Tris-HCl pH8.0, 0.01% Tween-20) containing 10% (w/v) bovine serum albumin (BSA) for 1 hour at room temperature before incubation overnight at 4°C with the indicated primary antibodies diluted 1 in 1000 with 5 % (w/v) BSA in TBS-T.

##### Antibodies

Mouse monoclonal anti-Myc (9E10, Santa Cruz Biotechnology), anti-HA (HA-7, Sigmaaldrich), anti-Flag or anti-Pk (V5 Tag) antibodies (Sigma-aldrich) were used for detecting epitope-tagged proteins. anti-phospho-P38 antibody (Thr180/Tyr182 Cell signalling) were used to detect the dual phosphorylated forms of Sty1 or P38. Total P38 and GAPDH from mammalian extracts were detected using rabbit derived antibody from Cell Signalling. Rabbit polyclonal anti-Hog1 antibodies (Y-215, Santa Cruz Biotechnology) or anti-Sty1 antibodies (Day and Veal 2010) were used to detect Sty1. Protein bands were visualised using the Licor Odyssey CLx system with Licor secondary antibody (IRDye 800 CW anti-rabbit/mouse) or anti-mouse IgG (whole molecule) (A4416) or anti-rabbit IgG (Sigma-Aldrich) conjugated to horseradish peroxidase.

##### Imaging and quantification of immunoblots

For Phos-tag^TM^ gels and experiments in Figures 2A, B, E and F; enhanced chemiluminescence (PierceTM ECL Plus, Thermo Scientific) and ImageQuant Software (Typhoon FLA9500) or X ray film (Fuji) was used to visualise detected proteins. All other immunoblots were imaged and quantified using Licor Image studio Lite: Lane normalization factor (LNF) was calculated from total protein stain (Revert ^TM^ LiCor) on 700 nm channel by selecting individual rectangles for appropriate equal-sized part of each lane after background subtraction in adjacent area (top and bottom/right and left). The signal of protein of interest was selected from 800nm channel with background subtraction. Normalised signal of target protein in each lane was then calculated by dividing the lane normalization factor for that lane. The exported value was then imported and plotted in Excel.

##### Determining *S. pombe* cell size at division

Cells that had been maintained continuously in exponential phase for 18-24 hrs were pelleted at OD _595_ 0.1 (Hagan 2016). Cells were fixed in 3.7% (w/v) formaldehyde solution for 10 min at room temperature, washed twice in phosphate buffered saline (PBS) (137 mM NaCl, 2.7 mM KCl, 10 mM Na_2_HPO_4_ and 1.8 mM KH_2_PO_4_) then resuspended in 10μl PBS solution containing 0.67 mg/ml Calcofluor (Sigma). Stained cells were mounted in Vectashield containing 1.5μg/ml 4, 6-diamidino-2-phenylindole (DAPI)(Vector Laboratory) on poly-L-lysine coated microscope slides. Cell morphology using differential interface contrast (DIC), and septum staining with Calcufluor, were imaged using Zeiss Axiovert or Axioskop fluorescence microscope (excitation wavelength 359nm and emission wavelength 461nm. The length of septum-stained cells was determined using Axiovison software for at least 150 cells per group.

##### Analysis of S. pombe cell volume using a Cell Counter

40μl of exponentially growing cells (OD595 0.3-0.5) were added to 10ml of CASYton solution, and following sonication, 3 repeated measurements of ~6,000 cells were made on the CASY Cell Counter + Analyser System, ModelTT (Schärfe System).

##### Experiments involving cells expressing analogue-sensitive Sty1^T97A^ mutants

SISA strains KS8266 and KS8311 (Mustavchiev et al., 2016) were maintained on Ye5S media containing 5μM 3-BrB-PP1(Abcam, ab143756) to prevent activation of sty1 and retention of Wis1^DD^ was confirmed by sequencing of cells used for experiments. For detecting Sty1 and Wis1 phosphorylation before and after oxidative stress, overnight SISA cells were grown in EMM media with supplements containing 5μM 3-BrB-PP1. Cells were diluted to OD_595_ 0.15 next day in the same media until reaching OD_595_ 0.5. Cell pellets were collected by <1 min centrifugation at < 2000rpm washed twice (one culture volume per wash) in either EMM media or EMM media containing 5 μM 3-BrB-PP1 before transfer to fresh EMM or EMM containing 5 μM 3-BrB-PP1 and treatment (where indicated) with 6 mM H_2_O_2_. To prepare cell pellets of control and sty1^T97A^ (MC138) strains, both cultures were grown in EMM with supplements without 3-BrB-PP1 overnight, followed by dilution next day until OD_595_=0.5. 5 μM 3-BrB-PP1 was then added, followed by 6 mM H_2_O_2_, and returned to incubator for a further 8 min before 10 min samples were collected (i.e. sty1^T97A^ kinase activity was inhibited for 10 min). Sty1 ^T97A^-Tpx1 expressing sty1 ^T97A^-tpx1 (MC146) and control cells were grown overnight in EMM then diluted to OD_595_= 0.15 in EMM containing (as indicated in figure) 5 μM 3BrB-PP1. After cells had grown to OD_595_=0.5 (~6h), cells were treated with H_2_O_2_ for 8 min (i.e. sty1^T97A^-tpx1 kinase activity was inhibited for ~6h).

### QUANTIFICATION AND STATISTICAL ANALYSIS

Each immunoblotting experiment shown in figures is representative of results from at least 3 independent biological repeats. Where genetic interactions were examined, results were repeated in multiple independent isolates in comparison with isogenic strains derived from the same cross. All experiments involving cell length measurements were repeated at least twice and results from representative experiments are shown. Error bars on graphs or in tables represent the standard error of the mean. As indicated, Student’s T tests were used to identify any quantitative differences between strains that were statistically significant and any differences p<0.05 are reported in figure legends.

## Supplemental Information

**Figure S1 Multiple cysteines in Sty1 are important for Sty1 activity and regulation by Tpx1 [A]** Immunoblot analysis (anti-Pk) of Δ*tpx1* mutant cells co-expressing Sty1-Pk (AD21) or Sty1^C35S^-Pk (EV59) with Tpx1^C170S^ (*Rep1Tpx1^C170S^*), Flag epitope-tagged Tpx1^C170S^ (*Rep1FlagTpx1^C170S^*) or vector control (*Rep1*) before and following 10min treatment with 0.2 mM H_2_O_2_ **[B]** Immunoblot analysis of activation of Sty1 by phosphorylation (anti-ppP38) before and following exposure of isogenic cells co-expressing wild-type Sty1 (AD38) or a Sty1^C35S^ mutant (AD40) and Tpx1 (*Rep1Tpx1*) or vector control (*Rep1*) to 0.2 mM H_2_O_2_ **[C]** Immunoblot analysis cells co-expressing wildtype Sty1 (AD38) or a Sty1^5CS^(Sty1^C13SC35SC153SC158SC242S^) mutant (AD84) in which 5 of the 6 cys are substituted with ser) with *tpx1^C170S^ (Rep1Tpx1^C170S^*) before and after 10 min exposure to 1 mM H_2_O_2_. Anti-Sty1 antibodies were used to detect Sty1. **[D]** DIC microscopy images of exponentially growing cells expressing wild-type (AD38) or *sty1^5CS^* (AD84). Pairs of newly divided cells are indicated by arrowheads. The mean length of septating *sty1*^+^ (AD38) and *sty1^5CS^* (AD84) cells as determined from images of calcufluor white-stained cells is indicated below each image. Error bars ± SEM, n=64 for each group shown, t-test, p=0.064 x 10^-21^. N=3 (a representative experiment is shown) **[E]** Mean volume of ~6000 wild-type (MG02) or *sty1^5CS^* (MG03) cells, as determined using a cell counter.**[F]** Immunoblot analysis of Sty1 phosphorylation (anti-ppP38) in cells expressing wild-type Sty1 (MC2), Sty1-tpx1 (MC12), Sty1^5CS^ (MC11) or Sty1-tpx1^C48S^ (MC82). Total Sty1 levels were determined using anti-Hog1 antibody. Total protein stain indicates protein loading. **[G]** Immunoblot analysis of Pyp1-Pk (anti-Pk) in cells expressing Pyp1-pk and wild-type Sty1(MC1), Sty1^5CS^ (MC5) or Sty1-Tpx1 (MC7) compared with tubulin loading control. In A and C the mobility of MW markers (kD) are indicated.

**Figure S2 FLAG-P38, FLAG-P38-Prdx1 and FLAG-P38-Prdx2 fusion proteins immunopurified from cells are active towards a model peptide substrate** Kinase assays were carried out using anti-Flag immunoprecipitates from cells expressing Flag-epitope-tagged P38, P38-Prdx1 or P38-Prdx2 and the fluorescent-tagged peptide substrate (5-FAM-IPTSPITTTYFFFKKK-COOH). The phosphorylation of the peptide substrate was ablated by inclusion of 10mM of the P38 inhibitor SB23903 in the reaction, confirming that the activity was due to immunopurified P38. Assays were repeated 3 times and results from a representative assay are shown. The presence of similar levels of FLAG-P38/FLAG-P38-Prdx fusion protein in assay were confirmed by immunoblotting.

**Figure S3 Tpx1 promotes Sty1 phosphorylation by mechanism/s that are independent of established mechanisms regulating Wis1 [A]** Overexpression of Tpx1 caused similar increases in H_2_O_2_–induced activation of Sty1 in cells that express HA-tagged wild-type Wis1 or Wis1 in which C458 is substituted with serine: Immunoblot analysis of Sty1 phosphorylation (anti-ppP38) in cells expressing wild-type Wis1(*wis1WT*;MC102) or Wis1^C458S^(*wis1^C458S^*;MC107) and over-expressing Tpx1 (from *Rep1tpx1^+^;* indicated ‘+’) compared with vector control (*Rep1;* indicated ‘-’) before and after 10 or 20 min exposure to 0.2 mM H_2_O_2_. Total Sty1 (anti-Hog1) and protein levels are indicated. **[B]** Immunoblot analysis of Sty1 phosphorylation (anti-ppP38) before and after exposure of cells coexpressing *wis1WT* and *wis1^C458S^* with wild-type Sty1 (MC110 and MC116), *sty1-tpx1* (MC112 and MC118) or *sty1-tpx1^C48S^* (MC114 and MC120) to H_2_O_2_ indicated that C458 in Wis1 is not required for the hyper-phosphorylation of Sty1-Tpx1 fusion proteins. MW markers are shown (kD) **[C-D]** Analysis of cells obtained from a cross between *wis1DD-myc* (NJ1088) and *sty1-tpx1* (MG17) expressing strains indicates that 4 different isolates bearing both alleles (MC84; i, ii, iii and iv) have adapted to the synthetic negative interaction by losing **[C]** Sty1-Tpx1 expression and **[D]** Sty1 activity, as indicated by the increased cell length of *‘sty1-tpx1 wis1DD’* (MC84) compared with wild-type cells (MC2). *wis1DD-myc* (MC19) and *sty1-tpx1* (MC12) expressing cells isogenic to MC84 are shown for comparison. Arrowheads indicate dividing cells.

**Figure S4 Analysis on Phos-tag™ gels reveals that Wis1 undergoes multiple phosphorylation events in response to oxidative and osmotic stress** Immunoblotting analysis with anti-myc antibodies of extracts from exponential phase cells expressing myc-epitope tagged Wis1 or Wis1^DD^ and separated on Phos-tag^TM^ gels. Cells were grown in Ye5S. **[A-C]** cells expressing myc-tagged wild-type Wis1 (KS2096) or Wis1^DD^ (KS2088) before and after exposure to the indicated concentrations of KCl (osmotic stress) or H_2_O_2_ **[D]** wild-type and Δ*mcs4* cells expressing Wis1myc before and following exposure to 6 mM H_2_O_2_ **[E-F]** wild-type and Δ*tpx1* cells expressing Wis1-myc or Wis1^DD^-myc before and following exposure to 6 mM H_2_O_2_ or 0.6 M KCl. In **A** duplicate samples are shown in which one has been treated with phosphatase to show that mobility shifts reflect phosphorylation of Wis1. Arrows indicate forms of Wis1 with different mobility. The number of ‘p’ preceding Wis1 is intended to reflect the extent of phosphorylation of different forms of Wis1.

**Figure S5: Sty1 kinase activity is not required for phosphorylation of Wis1. [A-B]** Wis1 undergoes a stress-induced mobility shift that does not require the MAP3K-phosphorylated residues in Wis1, Sty1 kinase, Pyp1 or Pyp2 activity. **[A]**The mobility of myc-tagged Wis1^DD^ was examined in Δ*pyp1Δpyp2* cells expressing an analogue sensitive Sty1^T97A^ mutant (KS8226) before and after 10 min exposure to 6mM H_2_O_2_. **[B]** Wis1 phosphorylation was investigated in Sty1(MC137) and Sty1^T97A^(MC138) cells expressing myc-tagged Wis1. Cells were grown in EMM media with appropriate supplements overnight then diluted and grown to OD_595_= 0.5. Cells were then treated with 5μM 3-BrB-PP1 and (as indicated) 6mM H_2_O_2_ for 10 min before collection. The H_2_O_2_ -induced mobility shift in Wis1 was detected in cells regardless of whether the Sty1 kinase activity was inhibited, indicating that Sty1 kinase activity is not required. **[C]** Cell expressing *sty1-tpx1* (MC12) or *sty1^T97A^-tpx1* (MC146) cells were grown in EMM media with supplements overnight. Cells were then diluted back to 0.15 next day in same media containing 5μM 3-BrB-PP1 until OD_595_ reached 0.5 (approximately 6 hours). The proportion of Wis1 that was phosphorylated (upper band) in Sty1^T97A^-Tpx1 expressing cells was similar regardless of the presence of the Sty1 kinase inhibitor (3-BrB-PP1).

**Figure S6: Effects of mutations in the MAPKK Wis1 on cell length: [A] Sty1-Tpx1^C48S^ fusion partially suppresses the cell cycle defect of cells expressing Wis1^AA^ (in which the canonical MAP3K sites are substituted with alanine) [B] Consistent with reduced Wis1 kinase activity and Sty1 activation (Fig. 6D), substitution of methionine 395 in Wis1 with alanine or glycine increases cell size at division** Sty1 activity is required for progression through G2 into mitosis [Millar et al., 1995; Shiozaki and Russell, 1995]. Cells from exponentially growing cultures co-expressing **[A]** wild-type Myc-tagged Wis1 and Sty1 (MG 132), Wis1^AA^-myc and wild-type Sty1 (MC122), Sty1-Tpx1^C48S^ and Wis1-myc (MC135), Wis1^AA^-myc and Sty1-Tpx1^C48S^ (MC130) or **[B]** wild-type Pk-tagged wild-type Wis1 (MG46), Wis1^M395A^ (MG47) or Wis1^M395G^(MG48) were imaged in comparison with Δ*sty1* (JM1160) mutant cells. Images were captured on [A] inverted microscope (Axiovert) or [B] using differential interference contrast (DIC) on upright microscope (Axioskop). Images shown were captured under the same conditions/magnification. Arrowheads indicate pairs of newly divided cells.

**Table S1 *S. pombe* strains used in this study** The construction of strains generated for this study is described in experimental procedures or below. *ade6* strains contain either *ade6-M210* or *ade6-M216*; where known, the particular allele is indicated. His^+^ strains in which it is possible but undetermined whether the *his7-366* allele is present are indicated by a question mark.

**Table S2 Primers used in this study**

**Table S3 Published plasmids used in this study**

