## Supplemental figures and information for "A Peroxiredoxin-P38 MAPK scaffold increases MAPK activity by MAP3K-independent mechanisms"

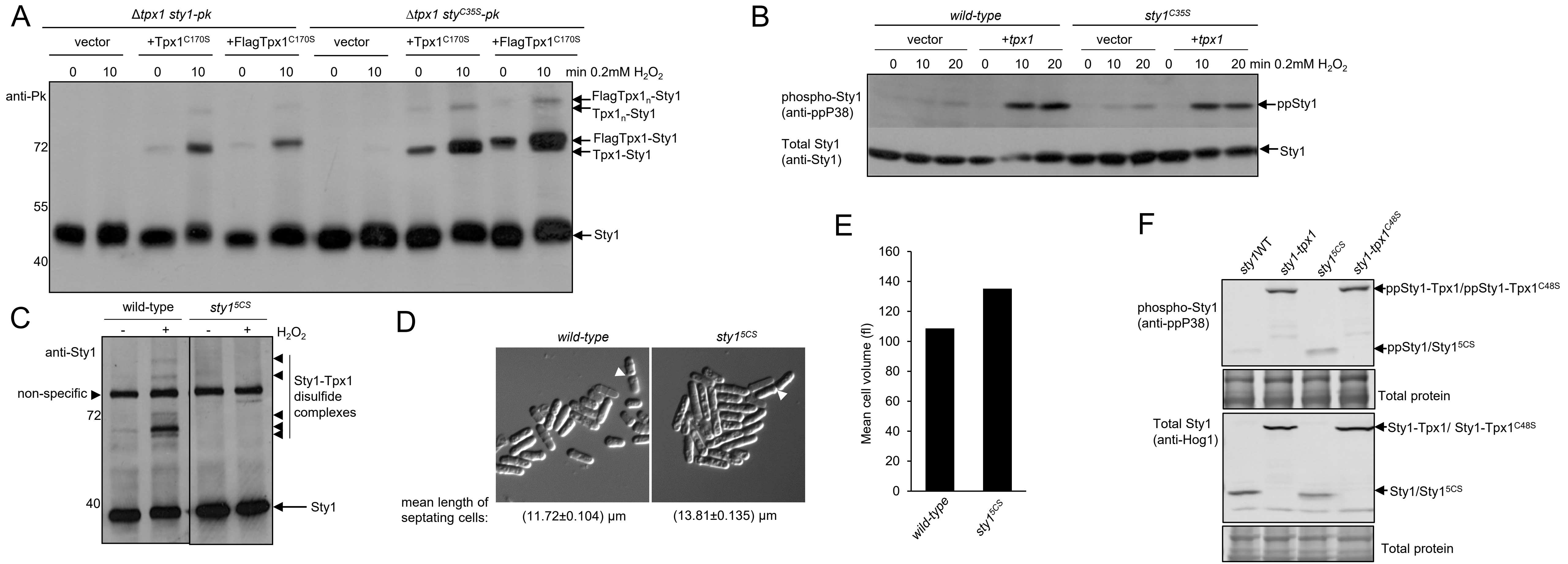


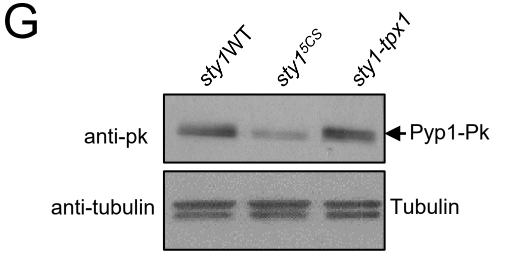


**Figure S1 Multiple cysteines in Sty1 are important for Sty1 activity and regulation by Tpx1 [A]** Immunoblot analysis (anti-Pk) of Δ*tpx1* mutant cells co-expressing Sty1-Pk (AD21) or $Sty1^{C35S}$-Pk (EV59) with $Tpx1^{C170S}$ (*Rep1Tpx1*$^{C170S}$), Flag epitope-tagged $Tpx1^{C170S}$ (*Rep1FlagTpx1*$^{C170S}$) or vector control (*Rep1*) before and following 10 min treatment with 0.2 mM $H_2O_2$ **[B]** Immunoblot analysis of activation of Sty1 by phosphorylation (anti-ppP38) before and following exposure of isogenic cells co-expressing wild-type Sty1 (AD38) or a $Sty1^{C35S}$ mutant (AD40) and Tpx1 (*Rep1Tpx1*) or vector control (*Rep1*) to 0.2 mM $H_2O_2$ **[C]** Immunoblot analysis of cells co-expressing wildtype Sty1 (AD38) or a $Sty1^{5CS}$($Sty1^{C13SC35SC153SC158SC242S}$) mutant (AD84) in which 5 of the 6 cys are substituted with ser) with *tpx1*$^{C170S}$ (*Rep1Tpx1*$^{C170S}$) before and after 10 min exposure to 1 mM $H_2O_2$. Anti-Sty1 antibodies were used to detect Sty1. **[D]** DIC microscopy images of exponentially growing cells expressing wild-type (AD38) or *sty1*$^{5CS}$ (AD84). Pairs of newly divided cells are indicated by arrowheads. The mean length of septating *sty1*$^{+}$ (AD38) and *sty1*$^{5CS}$ (AD84) cells as determined from images of calcufluor white-stained cells is indicated below each image. Error bars ± SEM, n=64 for each group shown, t-test, $p=0.064 \times 10^{-21}$. N=3 (a representative experiment is shown) **[E]** Mean volume of ~6000 wild-type (MG02) or *sty1*$^{5CS}$ (MG03) cells, as determined using a cell counter.**[F]** Immunoblot analysis of Sty1 phosphorylation (anti-ppP38) in cells expressing wild-type Sty1(MC2), Sty1-tpx1 (MC12), $Sty1^{5CS}$ (MC11) or Sty1-$tpx1^{C48S}$ (MC82). Total Sty1 levels were determined using anti-Hog1 antibody. Total protein stain indicates protein loading. **[G]** Immunoblot analysis of Pyp1-Pk (anti-Pk) in cells expressing Pyp1-pk and wild-type Sty1(MC1), $Sty1^{5CS}$ (MC5) or Sty1-Tpx1 (MC7) compared with tubulin loading control. In A and C the mobility of MW markers (kD) are indicated.

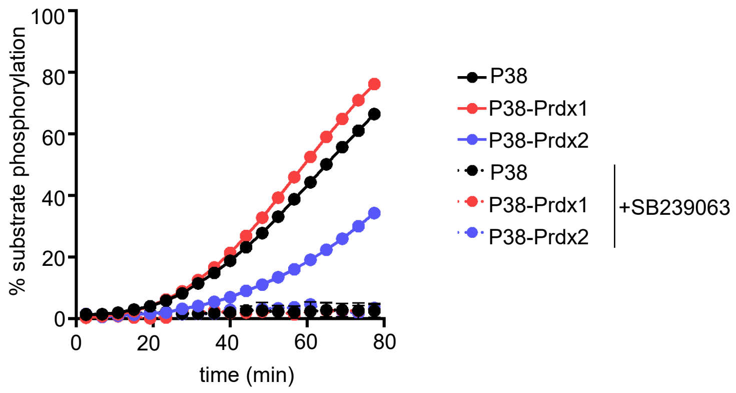


**Figure S2 FLAG-P38, FLAG-P38-Prdx1 and FLAG-P38-Prdx2 fusion proteins immunopurified from cells are active towards a model peptide substrate**
Kinase assays were carried out using anti-Flag immunoprecipitates from cells expressing Flag-epitope-tagged P38, P38-Prdx1 or P38-Prdx2 and the fluorescent-tagged peptide substrate (5-FAM-IPTSPITTTYFFFKKK-COOH). The phosphorylation of the peptide substrate was ablated by inclusion of 10μM of the P38 inhibitor SB23903 in the reaction, confirming that the activity was due to immunopurified P38. Assays were repeated 3 times and results from a representative assay are shown. The presence of similar levels of FLAG-P38/FLAG-P38-Prdx fusion protein in assay were confirmed by immunoblotting.

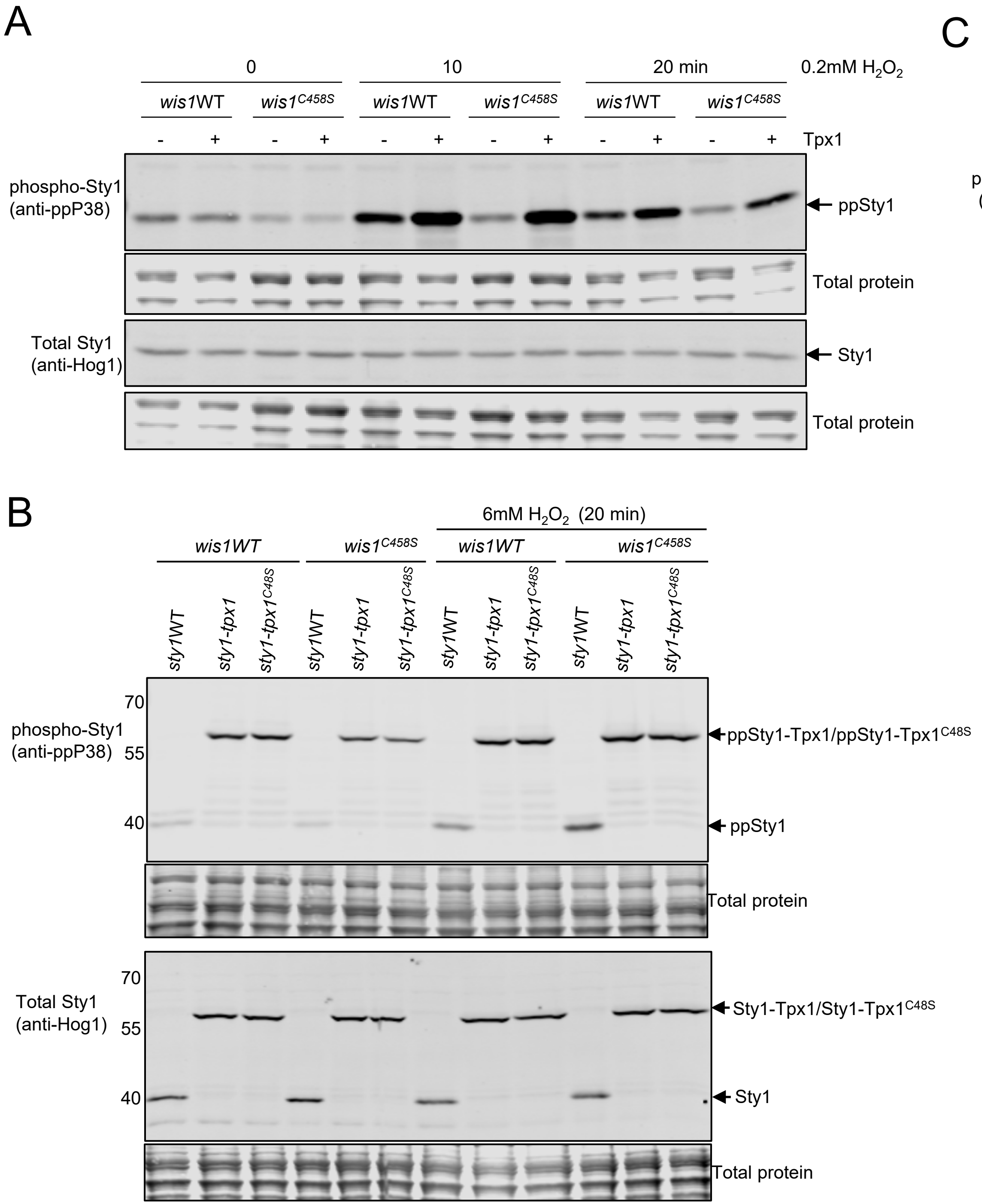


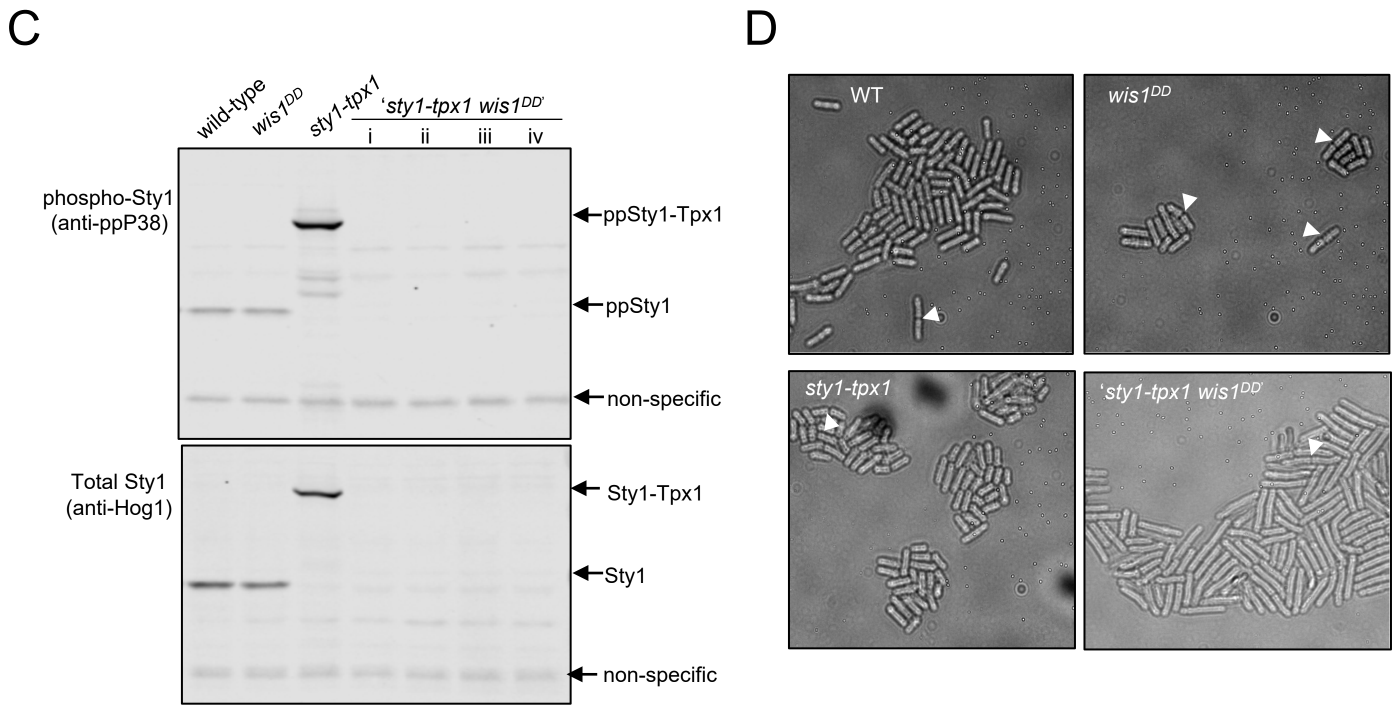


**Figure S3 Tpx1 promotes Sty1 phosphorylation by mechanism/s that are independent of established mechanisms regulating Wis1 [A]** Overexpression of Tpx1 caused similar increases in $H_2O_2$–induced activation of Sty1 in cells that express HA-tagged wild-type Wis1 or Wis1 in which C458 is substituted with serine: Immunoblot analysis of Sty1 phosphorylation (anti-ppP38) in cells expressing wild-type Wis1(*wis1WT*;MC102) or Wis1$^{C458S}$(*wis1*$^{C458S}$;MC107) and over-expressing Tpx1 (from *Rep1tpx1*$^{+}$; indicated '+') compared with vector control (*Rep1*; indicated '-') before and after 10 or 20 min exposure to 0.2 mM $H_2O_2$. Total Sty1(anti-Hog1) and protein levels are indicated. **[B]** Immunoblot analysis of Sty1 phosphorylation (anti-ppP38) before and after exposure of cells co-expressing *wis1*WT and *wis1*$^{C458S}$ with wild-type Sty1 (MC110 and MC116), *sty1-tpx1* (MC112 and MC118) or *sty1-tpx1*$^{C48S}$ (MC114 and MC120) to $H_2O_2$ indicated that C458 in Wis1 is not required for the hyper-phosphorylation of Sty1-Tpx1 fusion proteins. MW markers are shown (kD) **[C-D]** Analysis of cells obtained from a cross between *wis1*$^{DD}$*-myc* (NJ1088) and *sty1-tpx1* (MG17) expressing strains indicates that 4 different isolates bearing both alleles (MC84; i, ii, iii and iv) have adapted to the synthetic negative interaction by losing **[C]** Sty1-Tpx1 expression and **[D]** Sty1 activity, as indicated by the increased cell length of *'sty1-tpx1 wis1*$^{DD}$*'* (MC84) compared with wild-type cells (MC2). *wis1*$^{DD}$*-myc* (MC19) and *sty1-tpx1* (MC12) expressing cells isogenic to MC84 are shown for comparison. Arrowheads indicate dividing cells.

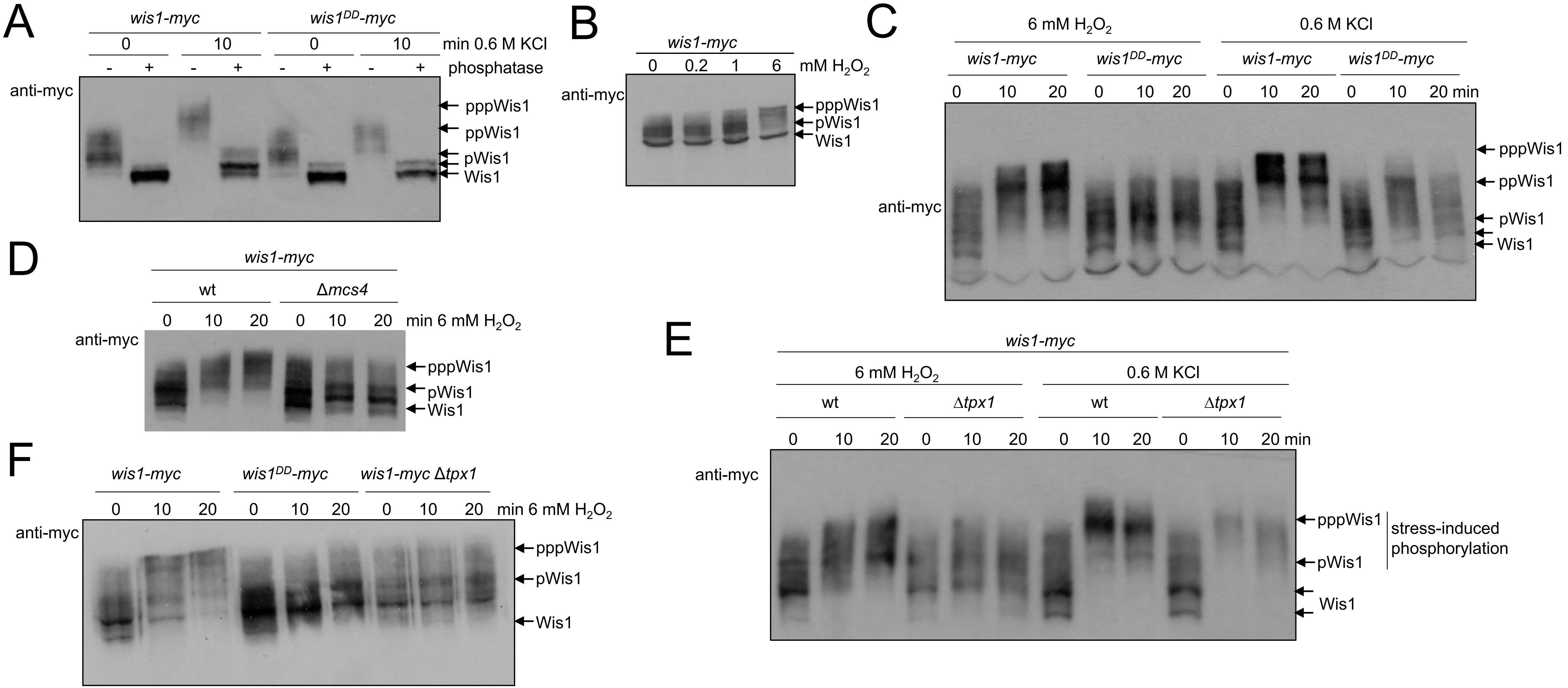


**Figure S4 Analysis on Phos-tag™ gels reveals that Wis1 undergoes multiple phosphorylation events in response to oxidative and osmotic stress** Immunoblotting analysis with anti-myc antibodies of extracts from exponential phase cells expressing myc-epitope tagged Wis1 or Wis1$^{DD}$ and separated on Phos-tag™ gels. Cells were grown in Ye5S. **[A-C]** cells expressing myc-tagged wild-type Wis1 (KS2096) or Wis1$^{DD}$ (KS2088**)** before and after exposure to the indicated concentrations of KCl (osmotic stress) or $H_2O_2$ **[D]** wild-type and Δ*mcs4* cells expressing Wis1-myc before and following exposure to 6 mM $H_2O_2$ **[E-F]** wild-type and Δ*tpx1* cells expressing Wis1-myc or Wis1$^{DD}$-myc before and following exposure to 6 mM $H_2O_2$ or 0.6 M KCl. In **A** duplicate samples are shown in which one has been treated with phosphatase to show that mobility shifts reflect phosphorylation of Wis1. Arrows indicate forms of Wis1 with different mobility. The number of ‘p’ preceding Wis1 is intended to reflect the extent of phosphorylation of different forms of Wis1.

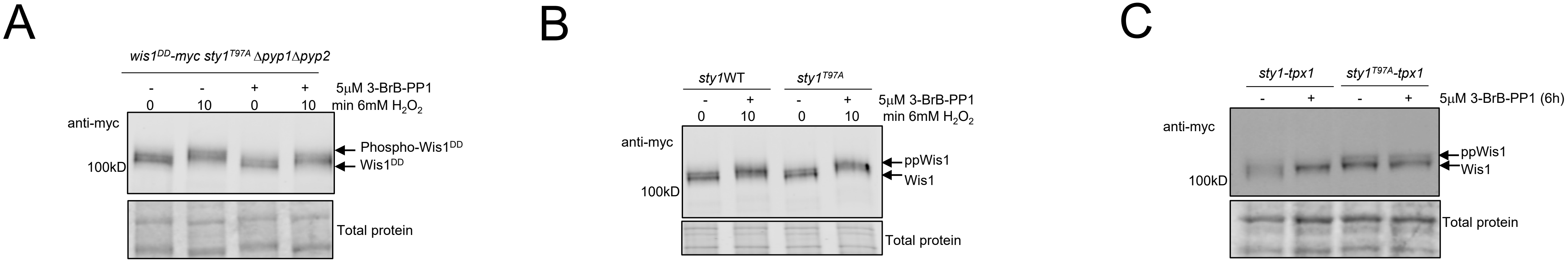


**Figure S5: Sty1 kinase activity is not required for phosphorylation of Wis1. [A-B]** Wis1 undergoes a stress-induced mobility shift that does not require the MAP3K-phosphorylated residues in Wis1, Sty1 kinase, Pyp1 or Pyp2 activity. **[A]** The mobility of myc-tagged Wis1$^{DD}$ was examined in Δ*pyp1*Δ*pyp2* cells expressing an analogue sensitive Sty1$^{T97A}$ mutant (KS8226) before and after 10 min exposure to 6 mM $H_2O_2$ . **[B]** Wis1 phosphorylation was investigated in Sty1(MC137) and Sty1$^{T97A}$(MC138) cells expressing myc-tagged Wis1. Cells were grown in EMM media with appropriate supplements overnight then diluted and grown to $OD_{595}$=0.5. Cells were then treated with 5μM 3-BrB-PP1 and (as indicated) 6 mM $H_2O_2$ for 10 min before collection. The $H_2O_2$ -induced mobility shift in Wis1 was detected in cells regardless of whether the Sty1 kinase activity was inhibited, indicating that Sty1 kinase activity is not required. **[C]** Cell expressing *sty1-tpx1* (MC12) or *sty1$^{T97A}$-tpx1* (MC146) cells were grown in EMM media with supplements overnight. Cells were then diluted back to 0.15 next day in same media containing 5μM 3-BrB-PP1 until $OD_{595}$ reached 0.5 (approximately 6 hours). The proportion of Wis1 that was phosphorylated (upper band) in Sty1$^{T97A}$-Tpx1 expressing cells was similar regardless of the presence of the Sty1 kinase inhibitor (3-BrB-PP1).

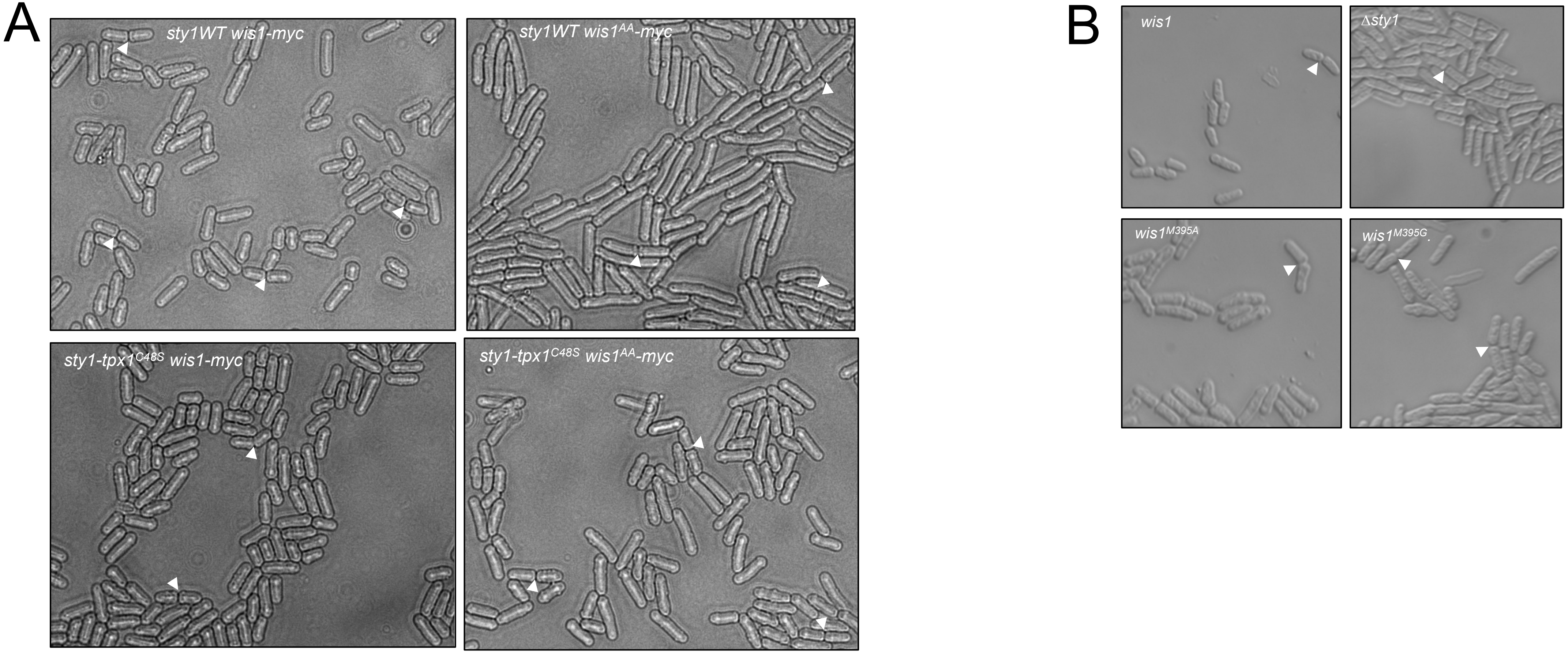


**Figure S6: Effects of mutations in the MAPKK Wis1 on cell length: [A] Sty1-Tpx1$^{C48S}$ fusion partially suppresses the cell cycle defect of cells expressing Wis1$^{AA}$ (in which the canonical MAP3K sites are substituted with alanine) [B] Consistent with reduced Wis1 kinase activity and Sty1 activation (Fig. 6D), substitution of methionine 395 in Wis1 with alanine or glycine increases cell size at division** Sty1 activity is required for progression through G2 into mitosis (Millar et al., 1995; Shiozaki and Russell, 1995). Cells from exponentially growing cultures co-expressing **[A]** wild-type Myc-tagged Wis1 and Sty1(MG132), Wis1$^{AA}$–myc and wild-type Sty1 (MC122), Sty1-Tpx1$^{C48S}$ and Wis1-myc (MC135), Wis1$^{AA}$-myc and Sty1-Tpx1$^{C48S}$ (MC130) or **[B]** wild-type Pk-tagged wild-type Wis1 (MG46), Wis1$^{M395A}$ (MG47) or Wis1$^{M395G}$(MG48) were imaged in comparison with Δ*sty1* (JM1160) mutant cells. Images were captured on [A] inverted microscope (Axiovert) or [B] using differential interference contrast (DIC) on upright microscope (Axioskop). Images shown were captured under the same conditions/magnification. Arrowheads indicate pairs of newly divided cells.

**Table S1 *S. pombe* strains used in this study** The construction of strains generated for use this study is described in experimental procedures or below. *ade6* strains contain either *ade6-M210* or *ade6-M216*; where known, the particular allele is indicated. His$^+$ strains in which it is possible but undetermined whether the *his7-366* allele is present are indicated by a question mark.

| strain | Genotype | Source (summary of construction details for unpublished strains) |
|---|---|---|
| AD13 | *h$^-$ ade6 leu1-32 his7-366 ura4-D18 sty1-3pk:ura4$^+$* | This study (pRip42PkSty1 linearised with BglII and integrated into CH429) |
| AD21 | *h$^-$ ade6 leu1-32 his7-366? ura4-D18 tpx1::ura4 sty1-3pk:ura4$^+$* | This study (Dissected spores from EV45 x AD13) |
| AD22 | *h$^-$ ade6-M216 his7-366 leu1-32 ura4-D18 sty1::his7$^+$* | Day and Veal (2010) |
| AD23 | *h- ade6-M216 leu1-32 ura4-D18 his7-366 sty1::his7$^+$ sty1$^{C35S}$-3pk:ura4$^+$* | This study (pRip42PkSty1$^{C35S}$ linearised with BglII and integrated into AD22) |
| AD38 | *h$^-$ ade6-M216 his7-366 leu1-32 ura4-D18 sty1::his7$^+$ sty1$^+$:ura4$^+$* | Day and Veal (2010) |
| AD40 | *h$^-$ ade6-M216 his7-366 leu1-32 ura4-D18 sty1::his7$^+$ sty1$^{C35S}$:ura4$^+$* | Day and Veal (2010) |
| AD84 | *h$^-$ ade6-M216 his7-366 leu1-32 ura4-D18 sty1::his7$^+$ sty1$^{C13SC35SC153SC158SC242S}$:ura4$^+$* | This study (pRip42Sty1$^{C13SC35SC153SC158SC242S}$ linearised with BglII and integrated into AD22) |
| AD130 | *h? trx1::kan$^{MX4}$ pyp1-3pk:kan $^{MX6}$ leu1-32 ura4-D18 his7-366 ade6-M210* | This study (Dissected spores from JB30 x AD142) |
| AD142 | *h- pyp1-3pk:kan$^{MX6}$ leu1-32 ura4-D18 his7-366 ade6-M210* | This study (NJ197 x CHP429) |
| AD143 | *h? ade6-M210 leu1-32 ura4-D18 pyp1-3pk:kan $^{MX6}$ tpx1::ura4$^+$* | This study (AD142 x VX00) |
| AD144 | *h? ade6-M210 his7-366 leu1-32 ura4-D18 Flag-trx1:ura4$^+$ trx1::kan$^{MX4}$ pyp1-3pk:kan$^{MX6}$* | This study (Dissected from JB35 x AD142) |
| CHP428 | *h$^+$ ade6-M210 leu1-32 ura4-D18 his7-366* | Laboratory stock |
| CHP429 | *h$^-$ ade6-M216 leu1-32 ura4-D18 his7-366* | Laboratory stock |
| EB15 | *h? ade6-M216 his7-366 leu1-32 ura4-D18 Leu1: nmt81:pyp2-13myc:ura4$^+$ pyp2::kan$^{MX6}$ sty1::his7$^+$ sty1$^+$:ura4$^+$* | This study: (Dissected from JP378 x AD38) |
| EB16 | *h? Leu1: nmt81:pyp2-13myc:ura4$^+$ pyp2::kan $^{MX4}$ sty1::his7$^+$ sty1-tpx1$^{C47S}$:ura4$^+$* | This study: (Dissected from JP378 x MG18) |
| EV45 | *h+ ade6 leu1-32 ura4-D18 tpx1::ura4$^+$* | Veal et al 2004 |
| EV59 | *h? ade6 leu1-32 ura4-D18 his7-366? sty1::his7$^+$ sty1$^{C35S}$-3pk:ura4$^+$ tpx1::ura4$^+$* | This study: (Dissected from EV45 x AD23) |
| HL2 | *h$^+$ ade6-M210 leu1-32 ura4-D18 pyp1-3pk:ura4$^+$* | This study (pRip42pyp1-3pk linearised and integrated into NT4) |
| JB30 | *h$^-$ trx1::kan$^{MX4}$ leu1-32 ura4-D18 his7-366 ade6-M210* | Brown et al 2013 |
| JJS7 | *h$^-$ kan$^{MX6}$:nmt1$^+$:3ha:wis1$^+$ his6:nat$^{MX6}$* | Sjölander et al 2020 (Gift of Prof Sunnerhagen) |
| JJS9 | *h$^-$ kan$^{MX6}$:nmt1$^+$:3ha:wis1$^{C458S}$ his6:nat$^{MX6}$* | Sjölander et al 2020 (Gift of Prof Sunnerhagen) |
| JM1160 | *h$^+$ ade6 his7-366 leu1–32 ura4-D18 sty1::ura4$^+$* | Millar et al 1995 (Gift of Prof Quinn) |
| JM1468 | *h$^+$ ade6-M210 his7-366 leu1-32 ura4-D18 mcs4::his7$^+$* | Buck et al, 2001 (Gift of Prof Quinn) |
| JP148 | *h$^+$ ade6-M216(0) his7-366 leu1-32 ura4-D18 wis1-12myc:ura4$^+$* | Gift of Dr.Quinn |
| JP279 | *h$^-$ pyp2::kan$^{MX6}$* | Petersen and Nurse, 2007 (Gift of Prof Petersen) |
| JP378 | *h$^+$ ura4-D18 leu1::nmt81.pyp2-13myc:ura4$^+$ pyp2::kan$^{MX6}$* | Petersen and Nurse, 2007 (Gift of Prof Petersen) |
| KS2086 | *h$^-$ leu1-32 ura4-D18 sty1:HA6H:ura4$^+$ wis1$^{AA}$-12myc:ura4$^+$* | Shiozaki et al. 1998 |
| KS2088 | *h$^-$ leu1-32 ura4-D18 sty1:HA6H:ura4$^+$ wis1$^{DD}$-12myc:ura4$^+$* | Shiozaki et al. 1998 |
| KS2096 | *h$^-$ leu1-32 ura4-D18 sty1:HA6H:ura4$^+$ wis1-12myc:ura4$^+$* | Shiozaki et al. 1998 |
| KS7830 | *h$^+$ sty1$^{T97A}$ ars1(BlpI):Padh13:CRIB-3xmCitrine:LEU2* | Mutavchiev et al., 2016 (Gift of Prof Sawin) |
| KS8226 | *h$^+$ leu1-32 ura4-D18 sty1$^{T97A}$ wis1$^{DD}$-12myc:ura4$^+$ pyp1::ura4$^+$ pyp2::LEU2 ars1(BlpI):Padh13:CRIB-3xmCitrine:LEU2* | Mutavchiev et al., 2016 (Gift of Prof Sawin) |
| KS8311 | *h$^+$ leu1-32 ura4-D18 sty1$^{T97A}$ wis1$^{DD}$-12myc:ura4$^+$ pyp1::ura4$^+$ pyp2::LEU2 ars1(BlpI):Padh13:CRIB-3xmCitrine:LEU2 Pact1:lifeactmCherry::leu1$^+$* | Mutavchiev et al., 2016 (Gift of Prof Sawin) |
| MC1 | *h$^-$ ade6-M216 his7-366 leu1-32 ura4-D18 pyp1-3pk kan $^{MX6}$ sty1::his7$^+$ sty1$^+$:ura4$^+$* | This study (Dissected from AD38 x NJ1088) |
| MC2 | *h$^-$ ade6-M216 his7-366 leu1-32 ura4-D18 pyp1-3pk:kan $^{MX6}$ sty1::his7$^+$ sty1$^+$:ura4$^+$ wis1-12myc:ura4$^+$* | This study (Dissected from MC1 x JP148) |
| MC5 | *h$^+$ ade6-M216 his7-366 leu1-32 ura4-D18 pyp1-3pk:kan $^{MX6}$ sty1::his7$^+$ sty1 $^{C13SC35SC153SC158SC242S}$:ura4$^+$* | This study (Dissected from AD84 x NJ1088) |

| | | |
|---|---|---|
| MC7 | h$^{-}$ ade6-M216 his7-366 leu1-32 ura4-D18 pyp1-3pk kan$^{MX6}$ sty1::his7$^{+}$ sty1-tpx1:ura4$^{+}$ | This study (Dissected from MG17 x NJ1088) |
| MC9 | h? ade6-M216 his7-366 leu1-32 ura4-D18 pyp1-3pk:kan$^{MX6}$ sty1::his7$^{+}$ sty1$^{C13SC35SC153SC158SC242S+}$-tpx1:ura4$^{+}$ | This study (Dissected from MG23 x NJ1088) |
| MC10 | h$^{-}$ his7-366 leu1-32 ura4-D18 wis1-12myc:ura4$^{+}$ | This study (Dissected from JP148 x CHP429) |
| MC11 | h? ade6-M216 his7-366 leu1-32 ura4-D18 pyp1-3pk kan$^{MX6}$ sty1::his7$^{+}$ sty1$^{C13SC35SC153SC158SC242S+}$:ura4$^{+}$ wis1-12myc:ura4$^{+}$ | This study (Dissected from MC10 x MC5) |
| MC12 | h$^{-}$ ade6-M216 his7-366 leu1-32 ura4-D18 pyp1-3pk kan$^{MX6}$ sty1::his7$^{+}$ sty1-tpx1:ura 4$^{+}$ wis1-12myc:ura4$^{+}$ | This study (Dissected from MC7 x JP148) |
| MC13 | h? ade6-M216 his7-366 leu1-32 ura4-D18 pyp1-3pk:kan$^{MX6}$ sty1:: his7$^{+}$ sty1$^{C13SC35SC153SC158SC242S+}$:ura4$^{+}$ -tpx1: ura4$^{+}$ wis1-12myc:ura4$^{+}$ | This study (Dissected from MC9 x JP148) |
| MC19 | ade6-216 his7-366 leu1-32 ura4-D18 pyp1-3pk kan$^{MX6}$ sty1::his7$^{+}$ sty1$^{+}$:ura 4$^{+}$ wis1$^{DD}$-12myc:ura4 | This study (Dissected from AD38 x NJ1088) |
| MC69 | h? ade6-M216 his7-366 leu1-32 ura4-D18 pyp1-3pk:kan$^{MX6}$ sty1::his7$^{+}$ sty1-tpx1$^{C48S}$:ura4$^{+}$ | This study (Dissected from MG18 x NJ1088) |
| MC77 | h? ade6-M216 his7-366 leu1-32 ura4-D18 pyp1-3pk:kan$^{MX6}$ sty1::his7$^{+}$ sty1:ura4$^{+}$ mcs4::his7$^{+}$ wis1-12myc:ura4$^{+}$ | This study (Dissected from MC2 x JP144) |
| MC82 | h$^{+}$ ade6-M216 his7-366 leu1-32 ura4-D18 pyp1-3pk:kan$^{MX6}$ sty1::his7$^{+}$ sty1-tpx1$^{C48S}$:ura4$^{+}$ wis1-12myc:ura4$^{+}$ | This study (Dissected from MC69 x JP148) |
| MC83 | h? ade6-M216 his7-366 leu1-32 ura4-D18 pyp1-3pk:kan$^{MX6}$ sty1::his7$^{+}$ sty1-tpx1:ura4$^{+}$ wis1-12myc:ura4$^{+}$ mcs4::his7$^{+}$ | This study (Dissected from MC12 x JP144) |
| MC84 | h? ade6-M216 his7-366 leu1-32 ura4-D18 pyp1-3pk:kan$^{MX6}$ sty1::his7$^{+}$ sty1-tpx1:ura4$^{+}$ wis1$^{DD}$-12myc:ura4$^{+}$ | This study (Dissected from MC7 x NJ1088 |
| MC98 | h? ade6-M216 his7-366 leu1-32 ura4-D18 sty1::his7$^{+}$ sty1$^{+}$:ura4$^{+}$ | This study (Dissected from CHP428 x AD38) |
| MC100 | h$^{+}$ ade6-M216 his7-366 leu1-32 ura4-D18 sty1::his7$^{+}$ sty1$^{+}$-tpx1$^{C48S}$:ura4$^{+}$ | This study (Dissected from CHP428 x MG18) |
| MC102 | h$^{+}$ ade6-M216 his7-366 leu1-32 ura4-D18 kan$^{MX6:}$nmt1$^{+}$:3ha:wis1$^{+}$ his6:nat$^{MX6}$ | This study (Dissected from CHP428 x JJS7) |
| MC103 | h$^{+}$ ade6-M216 his7-366 leu1-32 ura4-D18 kan$^{MX6}$:nmt1$^{+}$:3ha:wis1$^{+}$ his6:nat$^{MX6}$ | This study (Dissected from CHP428 x JJS7) |
| MC107 | h$^{+}$ ade6-M216 his7-366 leu1-32, ura4-D18 kan$^{MX6}$:nmt1$^{+}$:3ha::wis1$^{C458S}$ his6:Hph$^{MX6}$ | This study (Dissected from CHP428 x JJS9) |
| MC110 | h? ade6-M216 his7-366 leu1-32 ura4-D18 sty1::his7$^{+}$ sty1$^{+}$:ura4$^{+}$ Kan$^{MX6}$:nmt1$^{+}$:3ha:wis1$^{+}$ his6:nat$^{MX6}$ | This study (Dissected from MC102 x AD38) |
| MC112 | h? ade6-M216 his7-366 leu1-32 ura4-D18 sty1::his7$^{+}$ sty1-tpx1:ura4$^{+}$ kan$^{MX6}$:nmt1$^{+}$:3ha:wis1$^{+}$ his6:nat$^{MX6}$ | This study (Dissected from MC103 x MG17) |
| MC114 | h? ade6-M216 his7-366 leu1-32, ura4-D18 sty1::his7$^{+}$ sty1-tpx1$^{C48S}$:ura4+ kan$^{MX6}$:nmt1+:3ha:wis1+ his6:nat$^{MX6}$ | This study (Dissected from MC102 x MG18) |
| MC116 | h? ade6-M216 his7-366 leu1-32, ura4-D18 sty1::his7$^{+}$ sty1+:ura4+ kan$^{MX6}$:nmt1+:3ha:wis1$^{C458S}$ his6:Hph$^{MX6}$ | This study (Dissected from MC107 x AD38) |
| MC118 | h? ade6-M216 leu1-32, ura4-D18,his7-366, sty1::his7$^{+}$ sty1-tpx1:ura4+ kan$^{MX6}$:nmt1+:3ha:wis1$^{C458S}$ his6:Hph$^{MX6}$ | This study (Dissected from MC107 x MG17) |
| MC120 | h? ade6-M216 leu1-32, ura4-D18,his7-366, sty1::his7$^{+}$ sty1-tpx1$^{C48S}$:ura4+ kan$^{MX6}$:nmt1+:3ha:wis1$^{C458S}$ his6:Hph$^{MX6}$ | This study (Dissected from MC107 x MG18) |
| MC115 | h? ade6-M216 his7-366 leu1-32 ura4-D18 sty1::his7$^{+}$ sty1-tpx1$^{C48S}$:ura4$^{+}$ kan$^{MX6}$:nmt1$^{+}$:3ha:wis1$^{+}$ his6:nat$^{MX6}$ | This study (Dissected from MC102 x MG18) |
| MC122 | h? ade6-M216 his7-366 leu1-32 ura4-D18 sty1::his7$^{+}$ sty1$^{+}$:ura4$^{+}$ wis1$^{AA}$-12myc:ura4$^{+}$ | This study (Dissected from KS2086 x MC98) |
| MC130 | h? ade6-M216 his7-366 leu1-32 ura4-D18 sty1::his7$^{+}$ sty1-tpx1$^{C48S}$:ura4$^{+}$ wis1$^{AA}$-12myc:ura4$^{+}$ | This study (Dissected from KS2086 x MC100) |
| MC132 | h? ade6-M216 his7-366 leu1-32 ura4-D18 sty1::his7$^{+}$ sty1$^{+}$:ura4$^{+}$ wis1-12myc:ura4$^{+}$ | This study (Dissected from MC2 x MC98) |
| MC135 | h? ade6-M216 his7-366 leu1-32 ura4-D18 sty1::his7$^{+}$ sty1-tpx1$^{C48S}$:ura4$^{+}$ wis1-12myc:ura4$^{+}$ | This study (Dissected from MC82 x MG18) |
| MC137 | h? ade6-M216 sty1$^{+}$ wis1-12myc:ura4$^{+}$ | This study (Dissected from KS7830 x MC10) |
| MC138 | h? ade6-M216 sty1$^{T97A}$ wis1-12myc:ura4$^{+}$ | This study (Dissected from KS7830 x MC10) |
| MC146 | h? ade6-M216 his7-366 leu1-32 ura4-D18 pyp1-3pk:kan$^{1}$ sty1::his7$^{+}$ sty1$^{T97A}$-tpx1:ura4$^{+}$ wis1-12myc:ura4$^{+}$ | This study (pRip2Sty1$^{T97A}$-Tpx1 linearised and integrated into AD22, then x MC11) |
| MC150 | h? ade6-M216 his7-366 leu1-32 ura4-D18 sty1::his7$^{+}$ sty1-tpx1:ura4$^{+}$ pyp1::kan$^{MX6}$ | This study (Dissected from MG17 x NJ102) |
| MG02 | h$^{+}$ ade6-M216 his7-366 leu1-32 ura4-D18 sty1::his7$^{+}$ sty1$^{+}$:ura4$^{+}$ | This study (Dissected from AD38 x CHP428) |
| MG03 | h$^{+}$ ade6-M216 his7-366 leu1-32 ura4-D18 sty1::his7$^{+}$ sty1$^{C13SC35SC153SC158SC242S}$:ura4$^{+}$ | This study (Dissected from AD84 x CHP428) |
| MG17 | h$^{-}$ ade6-M216 his7-366 leu1-32 ura4-D18 sty1::his7$^{+}$ sty1-tpx1:ura4$^{+}$ | This study (pRip2Sty1-Tpx1 linearised and integrated into AD22) |
| MG18 | h$^{-}$ ade6-M216 his7-366 leu1-32, ura4-D18 sty1::his7$^{+}$ sty1-tpx1$^{C47S}$:ura4$^{+}$ | This study (pRip2Sty1-Tpx1$^{C47S}$ linearised and integrated into AD22) |
| MG23 | h$^{-}$ ade6-M216 his7-366 leu1-32 ura4-D18 sty1::his7$^{+}$ sty1$^{C13SC35SC153SC158SC242S}$-tpx1:ura4$^{+}$ | This study (pRip2Sty1$^{C13SC35SC153SC158SC242S}$-Tpx1 linearised and integrated into AD22) |
| MG46 | h$^{-}$ ade6-M216 leu1-32 ura4-D18 wis1-3pk:ura4$^{+}$ | This study (pRip42Wis1pkC linearised and integrated into NT4) |

| | | |
|---|---|---|
| MG47 | *h*$^{-}$ *ade6-M216 leu1-32 ura4-D18 wis1*$^{M395A}$*-3pk:ura4+* | This study (pRip42Wis1$^{M395A}$pkC linearised and integrated into NT4) |
| MG48 | *h*$^{-}$ *ade6-M216 leu1-32 ura4-D18 wis1*$^{M395G}$*-3pk:ura4*$^{+}$ | This study (pRip42Wis1$^{M395G}$pkC linearised and integrated into NT4) |
| MG49 | *h- ade6-M216 leu1-32 ura4-D18 sty1::his7*$^{+}$ *:sty1-tpx1:ura4*$^{+}$ *wis1-3pk:ura4*$^{+}$ | This study (Dissected from MG17 x MG46) |
| MG50 | *h*$^{-}$ *ade6-M216 leu1-32 ura4-D18 sty1::his7*$^{+}$ *:sty1-tpx1:ura4*$^{+}$ *wis1*$^{M395A}$*-3pk:ura4*$^{+}$ | This study (Dissected from MG17 x MG47) |
| MG51 | *h*$^{-}$ *ade6-M216 leu1-32 ura4-D18 sty1::his7*$^{+}$*:sty1-tpx1:ura4*$^{+}$ *wis1*$^{M395G}$*-3pk:ura4*$^{+}$ | This study (Dissected from MG17 x MG48) |
| NT4 | *h*$^{+}$ *ade6-M216 leu1-32 ura4-D18* | Laboratory stock |
| NT5 | *h*$^{-}$ *ade6-M216 leu1-32 ura4-D18* | Laboratory stock |
| NJ102 | *h*$^{+}$ *ade6-M210 his7-366 leu1-32 ura4-D18 pyp1::kan*$^{MX6}$ | Di et al 2011 (Gift of Drs C. Wilkinson and N. Jones) |
| NJ197 | *h*$^{+}$ *pyp1-3pk:kan*$^{MX6}$ *leu1-32 ura4-D18 his7-366 ade6-M210* | Gift of Drs C. Wilkinson and N. Jones |
| NJ1088 | *h*$^{+}$ *ade6-M216, leu1-32, ura4-D18 pyp1-3pk:kan* $^{MX6}$ *wis1*$^{DD}$*–12myc:ura4*$^{+}$ | Gift of Drs C Wilkinson and N. Jones |
| VX00 | *h*$^{+}$ *ade6 leu1-32 ura4-D18 tpx1::ura4*$^{+}$ | Veal et al 2004 |

### Table S2: List of primers used in this study

| primer | Oligonucleotide sequence 5'-3' |
|---|---|
| *pst1 sty1 mutant N* | CCCAGTCTGCAGCTCATGAACAACAATAAGGGAGTA |
| *sty1 internal F* | GAAGGATCAGGTAACTGG |
| *sty1 Nde1 R* | CGCCGCCATATGGGATTGCAGTTCATTATC |
| wis1_auto_*PstI*_Fwd | TATCTGCAGATGTCTTCTCCAAATAATCAACCC |
| wis1_auto_*KpnI*_Rev | ATAGGTACCCGCTTCTTTTTTCACCTTTCTCTTTA |
| wis1_mt395_chk_fwd | TGGCCTTGAAGGAAATTAGG |
| Pk_tag_check_rev | CAAGCAAAGGGTTAGGAATACCCAT |
| Pyp1intcheck | TTTAAGGCCAAATATTCTTAATAC |
| Pyp1intcheckB | TCTAAAATACCGAAGTCACCAGAA |

### Table S3 Published plasmids used in this study

| Plasmid | reference |
|---|---|
| pRep1 | Maundrell et al. (1988) |
| pRep1tpx1 | Veal et al., 2004 |
| pRep2tpx1 | Veal et al., 2004 |
| pRep2tpx1$^{C48S}$ | Veal et al., 2004 |
| pRip42pk | Day et al., 2012 |
| pcDNA3 | |
